# Single-cell multiomics data integration and generation with scPairing

**DOI:** 10.1101/2025.01.04.631299

**Authors:** Jeffrey Niu, Carlos Vasquez-Rios, Jiarui Ding

## Abstract

Single-cell multiomics technologies generate paired measurements of different cellular modalities, such as gene expression and chromatin accessibility. However, multiomics technologies are more expensive than their unimodal counterparts, resulting in smaller and fewer available multiomics datasets. Here, we present scPairing, a deep learning model inspired by Contrastive Language-Image Pre-Training, which embeds different modalities from the same single cells onto a common embedding space. We leverage the common embedding space to generate novel multiomics data following bridge integration, a method that uses an existing multiomics bridge to link unimodal data. Through extensive benchmarking, we show that scPairing constructs an embedding space that fully captures both coarse and fine biological structures. We then use scPairing to generate new multiomics data from retina, immune, and renal cells. Furthermore, we extend scPairing to generate trimodal data. Researchers can use these generated multiomics datasets to discover new biological relationships across modalities or confirm existing hypotheses.

## 1 Introduction

Recent advances in single-cell sequencing technologies have enabled the joint profiling of multiple modalities within individual cells.^1–4^ These multiomics technologies jointly profile different aspects of cell state, such as the transcriptome,^5^,^6^ chromatin accessibility,^7^ and surface epitopes.^4^ Single-cell multiomics facilitate a deeper characterization of cell state diversity and the interplay between different modalities,^8^,^9^potentially revealing the complex mechanisms that drive cellular processes.

Single-cell multiomics data integration presents a computational challenge due to the increased complexity inherent in such data. Computational methods must integrate the separate information sources in multiomics data to uncover patterns that are hard to detect from a single modality alone, such as gene regulatory relationships or unique cell states jointly defined by multiple modalities. To address these challenges, a wide range of single-cell multiomics integration tools have been developed to perform multiomics data integration. Some methods are specific to solely integrating paired data,^10–16^ while others are flexible in simultaneously integrating multimodal and unimodal data.^17–22^ Because of the scalability to handle large datasets and the effectiveness in capturing the structure in complex high-dimensional single-cell data, deep learning methods, especially deep generative models, are applied in multiomics integration methods.^10–13,15–18,21,22^ These models co-embed the multiomics data on a shared low-dimensional latent space. However, these models are typically more time-consuming to run compared to those for unimodal data analysis, and may rely on heuristics to combine the modality-specific embeddings to obtain joint cell-specific embeddings,^10^,^15,17,23^ or have difficulties embedding data in the presence of missing modalities.^12^,^14^,^16^Also, many of these models do not easily adapt to valuable multiomics data with more than two modalities.^24^,^25^

Despite the increasing availability and value of single-cell multiomics technologies, large, high-quality multiomics datasets remain limited due to high cost. A typical joint singlecell transcriptomics and epigenomics experiment usually produces fewer than 100,000 highquality cells, whereas unimodal sequencing using single-cell RNA sequencing (scRNA-seq) or single-cell assay for transposase-accessible chromatin with sequencing (scATAC-seq) has produced much larger data, including cell atlases with over a million cells.^26^,^27^ Multiomics data also have lower data quality compared to single-modality data, potentially hindering downstream analyses.^28^

To alleviate multiomics data quality and scarcity, many methods propose cross-modality imputation to smooth out noisy and sparse data and predict unobserved modalities from unimodal data. Many deep generative models include cross-modal decoders, which reconstruct expression profiles from the shared low-dimensional latent space. ^17^,^18^,^22^,^29^,^30^ However, these imputed data may miss fine-grained cell state information or have distorted characteristics compared to experimentally generated data. Consequently, these imputed data are seldom used for downstream analysis for biological discovery.

Here, we introduce scPairing, an efficient and flexible deep generative model for multiomics data integration and generation by pairing independently generated experimental unimodal data. scPairing has a modality-specific encoder to embed each modality data into a shared latent space. Furthermore, scPairing uses contrastive learning to encourage the modality-specific embeddings of the same cell to be nearby, while repelling embeddings of different cells. With contrastive learning alongside other objectives, scPairing prioritizes learning strongly aligned representations of individual modalities. This formulation enables both multiomics data integration and generation, where scPairing can produce joint embeddings suitable for biological investigation and generate artificial multiomics data by pairing independent experimental unimodal datasets, using a bridge multiomics dataset.^20^ By pairing unimodal data, our multiomics data generation procedure avoids over-smoothing seen by existing imputation methods. Furthermore, unlike other methods which all use highdimensional counts data as input, scPairing accepts low-dimensional embeddings for each modality, which enables efficient processing of atlas-level data.

We extensively test scPairing on fifteen datasets, demonstrating its superior performance on multimodal integration and pairing unimodal data. First, we demonstrate scPairing’s integration capabilities on common multiomics integration tasks. We then use the aligned embedding space learned by our model to perform cell pairing on human retina data, generating new multiomics profiles that we show to be more realistic than those produced by imputation methods. Second, we demonstrate the robustness of scPairing and the cell pairing procedure. Third, we apply scPairing to generate multiomics profiles in two distinct contexts: pairing a rare immune cell type across tissues, and pairing a clear cell renal carcinoma dataset We show how these artificially generated data can facilitate biological discovery. Lastly, we extend scPairing to integrate and generate single-cell trimodal sequencing data. Given its flexibility and scalability in processing large datasets, effectiveness in multiomics data integration, and its ability to pair large-scale unimodal data to generate multiomics data for biological discoveries, we envision that scPairing will become a valuable tool for single-cell multiomics data analysis. scPairing is a publicly available software tool, available at https://github.com/Ding-Group/scPairing.

## 2 Results

### scPairing jointly embeds cells onto a common hyperspherical space

The scPairing model uses a variational autoencoder (VAE), a deep generative model augmented with a contrastive loss, to embed single-cell multiomics data onto a common hyperspherical space (**Fig. 1A**).^31^ scPairing takes in low-dimensional representations of each modality computed from domain-specific methods such as principal component analysis (PCA), transcriptom specific VAEs,^31^,^32^or single-cell large language models (LLMs)^33^,^345^for scRNA-seq data, and latent semantic indexing (LSI) or PeakVI^35^ for scATAC-seq data. scPairing’s encoders then transform these low-dimensional representations onto a common hyperspherical latent space.

**Figure 1:**
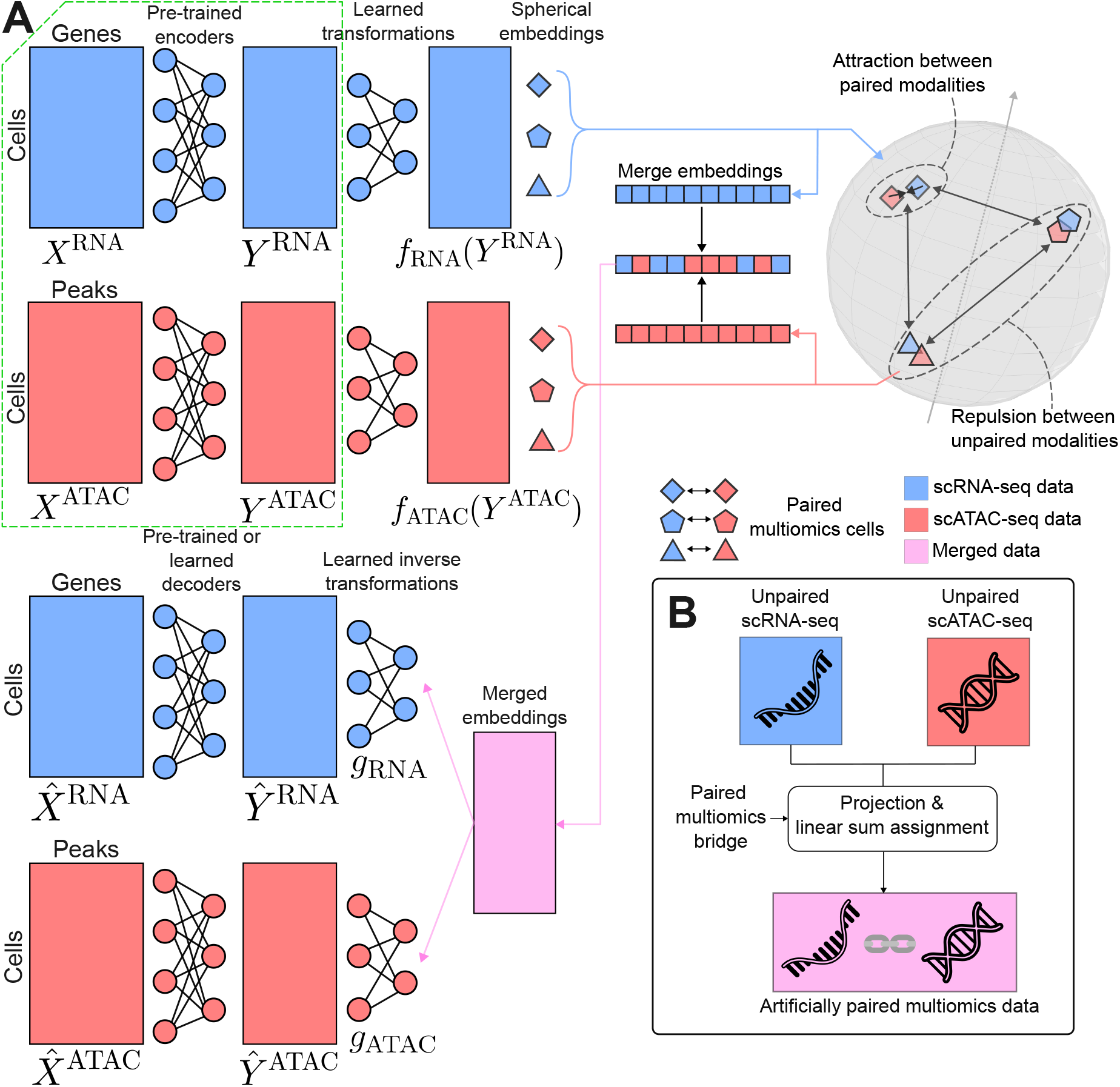
scPairing model overview. (A) Schematic of scPairing. Pre-trained encoders compute low-dimension embeddings *Y* ^RNA^ and *Y* ^ATAC^ that scPairing transforms onto a common hyperspherical embedding space. The dashed-green box indicates that the low-dimensional representations are assumed to be computed prior to running scPairing. scPairing applies a contrastive loss that encourages embeddings of two different modalities to embed close together, while repelling different cells’ embeddings. The two embeddings are merged and decoded back to their low-dimensional representations. Either a pre-trained decoder or a learned decoder reconstructs the count matrices for each modality. (B) Training scPairing on a multiomics bridge enables artificial pairing of independent scRNA-seq and scATAC-seq datasets. The scPairing encoders co-embed the two modalities in the same space. Solving the linear sum assignment problem with the similarities between scRNA-seq and scATAC-seq embeddings produces artificially paired data.

Following the Contrastive Language Image Pre-training (CLIP) approach,^36^ scPairing employs a contrastive objective that encourages transformed embeddings of different modalities from the same cell to be proximal, while embeddings from different cells are pushed apart. Our model adds further alignment objectives to ensure modality alignment (**Methods**).

Following the motivations of CLIP, we designed scPairing to take representations from modality-specific models by default. Using modality-specific representations provides flexibility in the final representations learned by scPairing, such as accounting for the trade-off between batch correction and biological conservation.^37^ Modality alignment becomes easier to learn with low-dimensional representations, as fewer dimensions make finding the alignment simpler than if the inputs were high-dimensional raw counts. Using low-dimensional representations also makes the application of scPairing computationally inexpensive, as scPairing has fewer parameters than a model that directly takes counts as input (**Supplementary Table S1**). The embeddings learned by scPairing can be used for downstream analyses, such as clustering and visualization.

Following a bridge integration procedure,^20^ scPairing can generate new multiomics data from separate unimodal measurements by artificially pairing unimodal cells (**Methods and Fig. 1B**). First, the individual modalities from the multiomics bridge data are integrated with the unimodal data. We train scPairing on only the bridge data to learn a common embedding space, then use its trained encoders to project the unimodal data into the same space We assign pairings between cells of the two modalities by computing the assignment of cells from one modality to the other modality such that the total similarity among all assignments is maximized (**Methods**).

### scPairing preserves biological structure while aligning modalities

We first applied scPairing to a joint scRNA-seq and scATAC-seq benchmarking dataset of bone marrow mononuclear cells (BMMCs).^38^ For the scRNA-seq low-dimensional representations, we applied embeddings from Harmony-corrected PCA,^39^ scVI,^32^ scPhere,^31^ scGPT,^33^ and CellPLM,^34^ while for the scATAC-seq low-dimensional representations, we applied embeddings from Harmony-corrected LSI and PeakVI.^35^

To quantify the preservation of biological structure, we computed cross-batch *k*-nearest neighbor cell type classification accuracies with scPairing embeddings. We held out cells from one batch and predicted their types using the cells from the other batches, which penalizes embeddings that fail to remove batch effects. Our results show scPairing preserves biological structure just as well as other multiomics methods that learn the structure from count data, and better than unimodal methods (**Fig. 2A**). We also performed ten-fold crossvalidation of *k*-nearest neighbors accuracies, where we saw similar high performance from scPairing (**Fig. 2A**). These results demonstrate the tradeoff between batch correction and biological conservation, as scVI and scPhere scRNA-seq embeddings for scPairing performed best in cross-batch classification, while CellPLM scRNA-seq embeddings performed best in ten-fold cross-validation classification.

**Figure 2:**
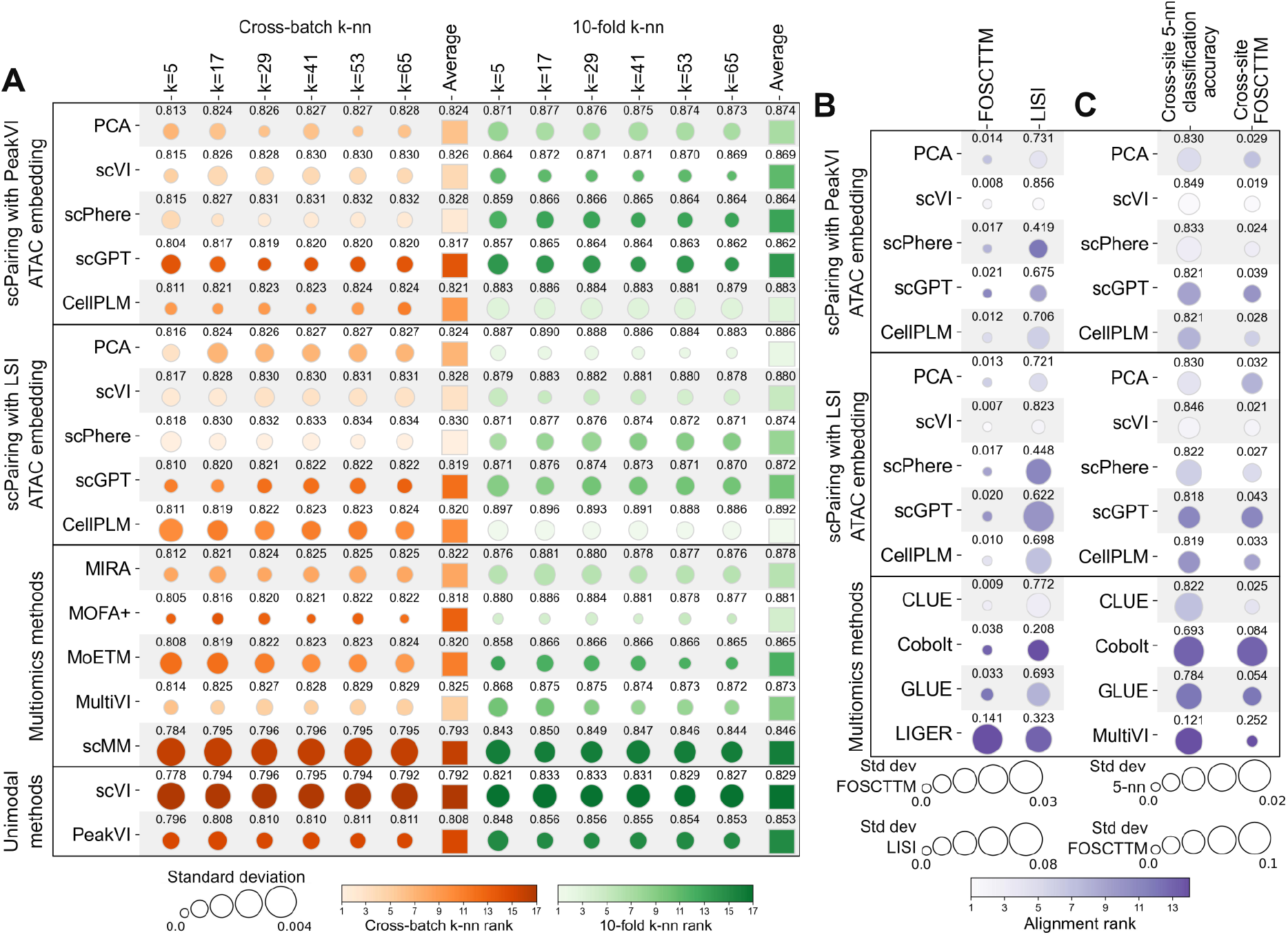
Benchmarking integration of joint scRNA-seq and scATAC-seq data. (A) Comparison of biological structure preservation quantified by *k*-nearest neighbor cell type classification accuracies. We predicted cell types for one batch given the cells from the remaining batches (cross-batch *k*-nn) or predicted cell types across 10-fold cross-validation (10-fold *k*-nn). (B) Comparison of modality alignment performance with FOSCTTM and LISI computed across all cells. (C) Comparison of bridge integration performance. Cross-site 5-nn classification accuracy assesses cell type classification using the held-out scRNA-seq profiles as a reference to predict the scATAC-seq cell types. Cross-site FOSCTTM assesses the alignment of the heldout scRNA-seq and scATAC-seq profiles. The row labels for scPairing in A–C are the scRNA-seq modality representation used as input. Each of experiments in A and B were repeated for five trials. Each of the experiments in C was repeated for five trials per held-out site. The mean of each metric is labeled and the circle size denotes the standard deviation.

Next, we evaluated scPairing’s ability to align the two modalities in the common embedding space using the Fraction of Samples Closer than True Match (FOSCTTM) metric (**Methods**).^40^ We observed that scPairing embeddings derived from scVI with PeakVI or Harmony-corrected LSI outperformed four other methods designed for modality alignment, including CLUE, Cobolt, GLUE, and LIGER (**Fig. 2B**).

Multiomics alignment is important as it imbues each modality’s encoder with rich information from other modalities, such as differentiation patterns or modality-specific cell types. We observed this transfer of information when applying scPairing to data with modality-specific information. First, we used a dataset of mouse skin cells sequenced by SHAREseq,^2^ with Palantir^41^ used to order the cells in pseudotime. Initially, only the scATAC-seq data showed the correct differentiation trajectory.(**Supplementary Fig. S1A and S1B**). After applying scPairing, both the scRNA-seq and scATAC-seq embeddings followed the correct differentiation trajectory. (**Supplementary Fig. S1C and S1D**). Second, we applied scPairing to a set of joint scRNA-seq and scATAC-seq cells from the developing human cerebral cortex.^42^ In this data, three blood vessel cell subtypes (endothelial cells, pericytes, and vascular smooth muscle cells) share similar chromatin accessibility profiles, but have distinct transcriptomes (**Supplementary Fig. S1E and S1F**). After aligning the modalities, the embeddings for the scATAC-seq modality show a clearer distinction between the three cell subtypes (**Supplementary Fig. S1G and S1H**). We confirmed these observations by computing the per-cell type Local Inverse Simpson’s Index (LISI), which showed a sharp increase in all cell types in the scATAC-seq modality, while remaining mostly unchanged in the scRNA-seq modality (**Supplementary Fig. S1I and S1J**). Focusing specifically on the separation of the three blood vessel cell subtypes, we observed a similar trend (**Supplementary Fig. S1K and S1L**).

We also investigated modality mixing in scPairing’s representations, as aligning the modalities does not preclude the possibility that the modalities are separated in the common embedding space. We calculated the LISI, which measures the diversity within the nearest neighborhoods in a *k*-nearest neighbor graph. We saw that scPairing embeds modalities with greater mixing compared to four other modality alignment methods (**Fig. 2B**).

Next, we compared scPairing to other methods in bridge integration, which is key to achieving high-quality cell pairings. To simulate bridge integration, we held out one of four sequencing sites from the benchmarking data, using three sequencing sites as the bridge data, and the held-out site as two separate unimodal data. We trained each method on the bridge data and evaluated their ability to classify the held-out scATAC-seq cell types using the held-out scRNA-seq cell type labels as a reference, and their alignment of the two modalities from the held-out data (**Methods**). In the cell type classification task, we observed that scPairing is competitive with other methods, while in the alignment task, scPairing better aligns modalities than other methods (**Fig. 2C**).

Though we argue for cell pairing for multiomics data generation, we also tested scPairing in performing cross-modal imputation since both decoders in scPairing decode from the common latent space. We performed the same cross-site split of the benchmarking dataset and used the scATAC-seq embeddings to impute scRNA-seq profiles and vice-versa. scPairing with pre-trained or learned decoders achieved comparable imputations based on the Spearman correlation between true and imputed gene expression (**Supplementary Fig. S2A**), and the imputations accurately reflected the expression of marker genes (**Supplementary Fig. S2B and S2C**). The binary cross-entropy between the true binarized and imputed peak accessibility was lower in three of four sites when comparing scPairing with pre-trained decoders against MultiVI or scButterfly (**Supplementary Fig. S2D**). When using learned decoders, the binary cross-entropy drastically decreased, with performance comparable to BABEL.

Lastly, we demonstrate scPairing’s flexibility to learn aligned representations of CITE-seq data, another multiomics technology that jointly measures gene expression and cell surface proteins.^4^ We applied scPairing to two CITE-seq datasets: 90,261 BMMCs from Luecken *et al*.^38^ *and 161,764 peripheral blood mononuclear cells (PBMCs) from Hao et al*.^14^ *First, we evaluated biological structure preservation with cross-batch k*-nearest neighbor classification accuracies. We observed that scPairing achieved the best accuracy in the BMMC data and the second-best accuracy in the PBMC data (**Supplementary Fig. S3A and S3B**). Second, we evaluated modality alignment with FOSCTTM and LISI, where we observed that scPairing was the only method to achieve both low FOSCTTM and high LISI in both datasets (**Supplementary Fig. S3C**). While CLUE achieved strong FOSCTTM, it failed at mixing the modalities, and while LIGER achieved strong mixing, it failed to align modalities originating from the same cell. These results show scPairing’s flexibility in integrating multiomics datasets across various modalities and technologies, achieving superior performance over standard methods.

### Bridge integration produces artificial cell pairings resembling true multiomics data

We evaluated scPairing’s ability to produce realistic artificial multiomics data by pairing gene expression and chromatin accessibility profiles from human retinal cells, using another paired multiomics dataset as a bridge. The unpaired Liang *et al*. dataset contains over 250,000 cells sequenced with single-nuclei RNA sequencing (snRNA-seq) and 137,000 cells sequenced with single-nuclei ATAC-seq (snATAC-seq).^43^ The paired Wang *et al*. bridge dataset contains over 45,835 joint scRNA-seq and scATAC-seq cells.^44^

After harmonizing the unpaired and bridge datasets and applying scPairing, we obtained 105,613 artificially paired cells (**Fig. 3A**). Visualization of our artificial pairings shows that scPairing correctly separated the discrete major cell types in the retina. To determine the quality of our pairings, we focused on 24,446 retinal bipolar cells. Liang *et al*. annotated 14 retinal bipolar cell subtypes, which our pairings also successfully captured (**Fig. 3B**).

**Figure 3:**
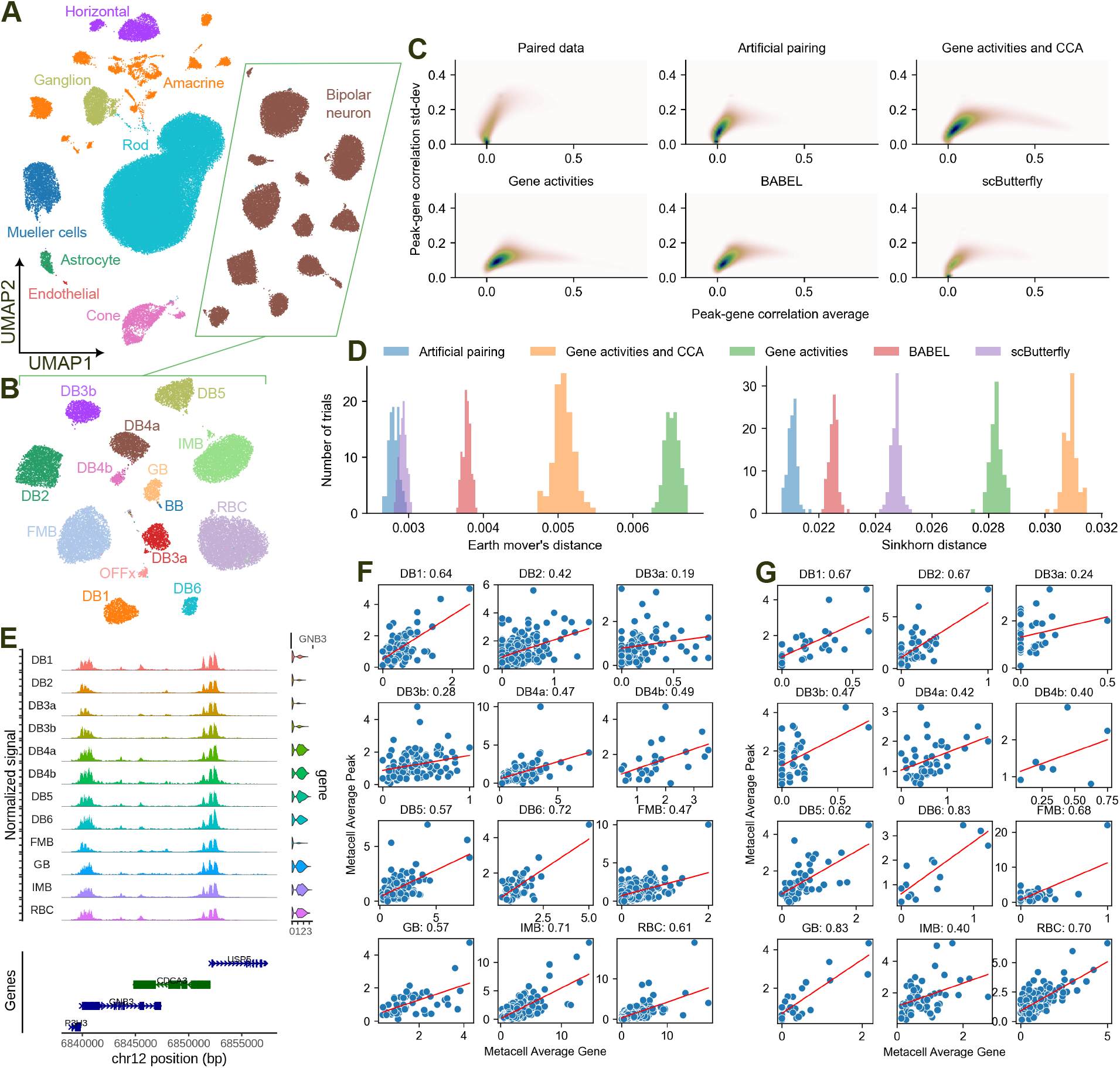
Artificial pairing of human retinal data. (A) Uniform manifold approximation and projection (UMAP) visualization of the artificially paired human retinal cells produced by scPairing, colored by major cell type. (B) UMAP visualization of the retinal bipolar subset from the artificial pairing, colored by cell subtype. (C) Comparison of density plots of peak-gene correlation means and standard deviations across the retinal bipolar subtypes on the Wang *et al*. multiomics data ^44^ (top-left), scPairing artificial pairings of the Liang *et al*. unpaired data ^43^ (top-middle), imputed gene expression of the Liang *et al*. scATAC-seq data using gene activities followed by CCA with scRNA-seq data (top-right), using only gene activities (bottom-left), BABEL (bottom-middle), and scButterfly (bottom-right). Correlations were computed using metacells produced by SEACells. ^48^ (D) Earth Mover and Sinkhorn distances between the distribution of gene-peak correlation means and standard deviations for the Liang *et al*. dataset computed from the different multiomics data generation methods compared to the distribution from the Wang *et al*. dataset. (E) Coverage plot of retinal bipolar subtypes at the *GNB3* TSS. (F and G) Correlation of average *GNB3* gene expression and chr12:6850801-6853197 accessibility across 12 retinal bipolar subtypes aggregated into metacells from our artificial pairings (F) and the Wang *et al*. multiomics data (G).

We further assessed whether our artificial pairings are concordant with true paired data by calculating the correlation between gene expression and chromatin accessibility of peaks within 250kb of the gene transcription start site (TSS) across 12 abundant retinal bipolar cell subtypes. We compared our pairings against imputations from canonical correlation analysis (CCA) between gene activities and the scRNA-seq data,^45^ imputations solely derived from gene activities,^46^ and two deep learning methods, BABEL^29^ and scButterfly.^47^ Sparsity presents a challenge in computing faithful gene-peak correlations. Thus, we applied the SEACells algorithm^48^ to aggregate retinal bipolar cell subtypes into robust meta-cells (**Methods**). For each gene-peak pair, we computed the mean and standard deviation of the Pearson correlations across the 12 subtypes (**Fig. 3C**).

We found that the gene-peak correlations of the artificial pairings most closely match the correlations from the paired multiomics data (**Fig. 3D**). The multiomics data suggest weak correlations between most genes and peaks close to the TSS have little correlation with small variance between subtypes. Artificial pairings produced by scPairing have a distribution of means and standard deviations closest to the paired data. The correlations produced by gene activities, gene activities followed by CCA, and BABEL all overestimate the gene-peak correlations and variance of correlations between subtypes. For gene activities, the direct transformation from gene accessibility likely neglects the variability present within cells, while neural networks tend to oversmooth when making predictions. scButterfly manages to produce a distribution closer to the paired data, notably by having a substantial number of uncorrelated gene-peak pairs.

We then re-analyzed the relationship between *GNB3* expression and the peak at chr12:685080 6853197 (**Fig. 3E**). Using aggregated metacells, we found that this gene-peak pair has a strong positive correlation across all subtypes (**Fig. 3F**). This result aligns with the correlations seen in the paired multiomics data, which also showed positive correlations (**Fig. 3G**). In contrast, the original study aggregated cells by donor, which produced heterogeneous correlations because cells from different donors were similar (**Methods, Supplementary Fig. S4**)

### Robustness of scPairing

We performed ablation studies on the scPairing bridge integration procedure to evaluate its robustness in data integration and pairing. First, we split the benchmarking BMMC dataset into a bridge dataset consisting of three of the four sites, and a test dataset consisting of one site separated into its individual modalities for alignment and pairing. We downsampled each cell type in the bridge dataset to 2%, 1%, 0.1% and 0% of the total number of cells to investigate how scPairing handles rare cell types. We observed that most cell types’ FOSCTTM did not drastically increase as the cell type was downsampled in the bridge dataset (**Fig. 4A**). We then assessed whether scPairing could correctly re-pair cells of the downsampled cell types in the test dataset. We observed that, with the exception of pDCs, all cell types retained a similar proportion of cell type-consistent re-paired cells (**Methods and Fig. 4B**).

**Figure 4:**
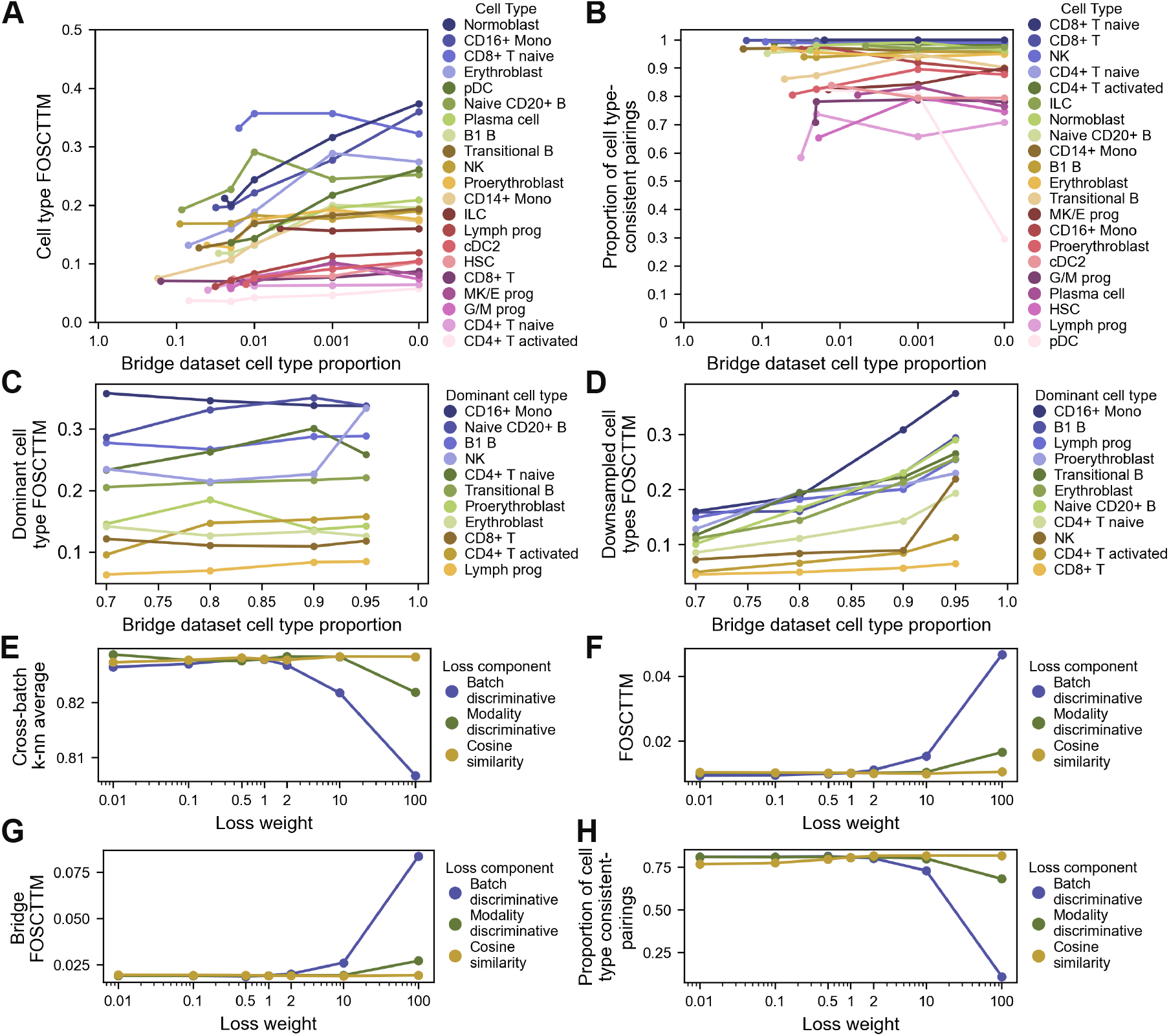
Robustness and sensitivity of scPairing and artificial pairing. (A and B) Robustness of bridge integration and pairing in the presence of rare cell types in the bridge data. We assessed modality alignment using FOSCTTM (A) and artificial pairing validity using the proportion of cell type-consistent pairings (B). The first point for each line represents the original cell type proportion in the bridge data. (C and D) Robustness of bridge integration in the presence of a dominant cell type. We assessed the modality alignment of the dominant cell type (C) and the downsampled cell types (D) using FOSCTTM. (E–H) Sensitivity of scPairing’s additional losses in multiomics integration (E and F), bridge integration (G), and pairing (H). We assessed scPairing’s learned embeddings in multiomics integration using cross-batch *k*-nearest neighbor classification accuracy (E) and alignment using FOSCTTM (F). Bridge integration was assessed using the FOSCTTM of the held out site (G). Pairing was assessed using the proportion of cell type-consistent pairings (H).

Next, we downsampled all but one cell type in the bridge dataset such that one cell type became dominant, comprising 70%, 80%, 90%, or 95% of the cells in the bridge. After applying scPairing to embed the test data onto the common embedding space, the FOSCTTM for the dominant cell type remained stable, while the rest of the cell types only exhibited a marked increase in FOSCTTM when the dominant cell type proportion increased from 90% to 95% (**Fig. 4C and 4D**). Altogether, these results suggest that modality alignment and cell pairing with scPairing is robust to variations in cell composition in the bridge data, notably learning good alignments on underrepresented cell types.

The scPairing model consists of three additional losses: a modality discriminative loss, a batch discriminative loss, and a cosine similarity alignment loss. We evaluated scPairing with all three combinations of the losses being either enabled or disabled. While we did not observe drastic differences in the FOSCTTM across the eight model variations, we noticed that including the cosine similarity alignment loss increased the number of paired cells (**Supplementary Table S2**). The effects of the three losses became more apparent in the retinal experiment, where the highest proportion of cells were paired when applying scPairing with all three losses enabled (**Supplementary Table S2**).

We also tested the sensitivity of the three additional losses by increasing or decreasing their weight in the overall scPairing loss (**Fig. 4E–H**). We observed that the cosine similarity loss weight does not affect the performance in multiomics integration or bridge integration and pairing, while the batch discriminative and modality discriminative losses only deteriorate performance at high weights, suggesting that the current loss weights lead to good performance.

Lastly, we modified the *ϵ* parameter used during pairing to observe how the yield of paired cells varies. In both the test dataset and the retinal experiment, the number of cells paired only drops drastically when *ε* decreases past 0.05 (**Supplementary Table S3**). However, the effect of this parameter is data-dependent and should be adjusted according to the desired similarity between paired profiles.

### Using a specialized bridge dataset pairs a rare immune cell type

Ulezko Antonova *et al*. recently described a new rare cell type with characteristics of both dendritic cells (DCs) and type 3 innate lymphoid cells (ILC3s), which they termed ROR*γ*t^+^-DC (R-DC-like) cells.^49^,^50^ In their study, they applied a sorting strategy to yield a joint scRNA-seq and scATAC-seq dataset of human tonsil cells enriched for R-DC-like cells. To test the versatility of scPairing, we attempted to pair scRNA-seq and scATAC-seq data from two datasets where very few R-DC-like cells have been captured, using the Ulezko Antonova *et al*. dataset as a bridge. To evaluate the quality of our pairings, we identified cells that express *PRDM16, RORC, PIGR*, and *FOXP2* as candidate R-DC-like cells. If we successfully paired these cells, the artificially paired cells should show a chromatin accessibility profile similar to the R-DC-like cells in the Ulezko Antonova *et al*. dataset (**Fig. 5A**). We highlight four regions examined by Ulezko Antonova *et al*. at the *RORC, AIRE*, and *CD4* loci. They found that R-DC-like cells have an accessible *RORγt* promoter region but not at the *RORγ* promoter region, an inaccessible *AIRE* conserved non-coding sequence 1 (CNS1), and an accessible *CD4* promoter region.

**Figure 5:**
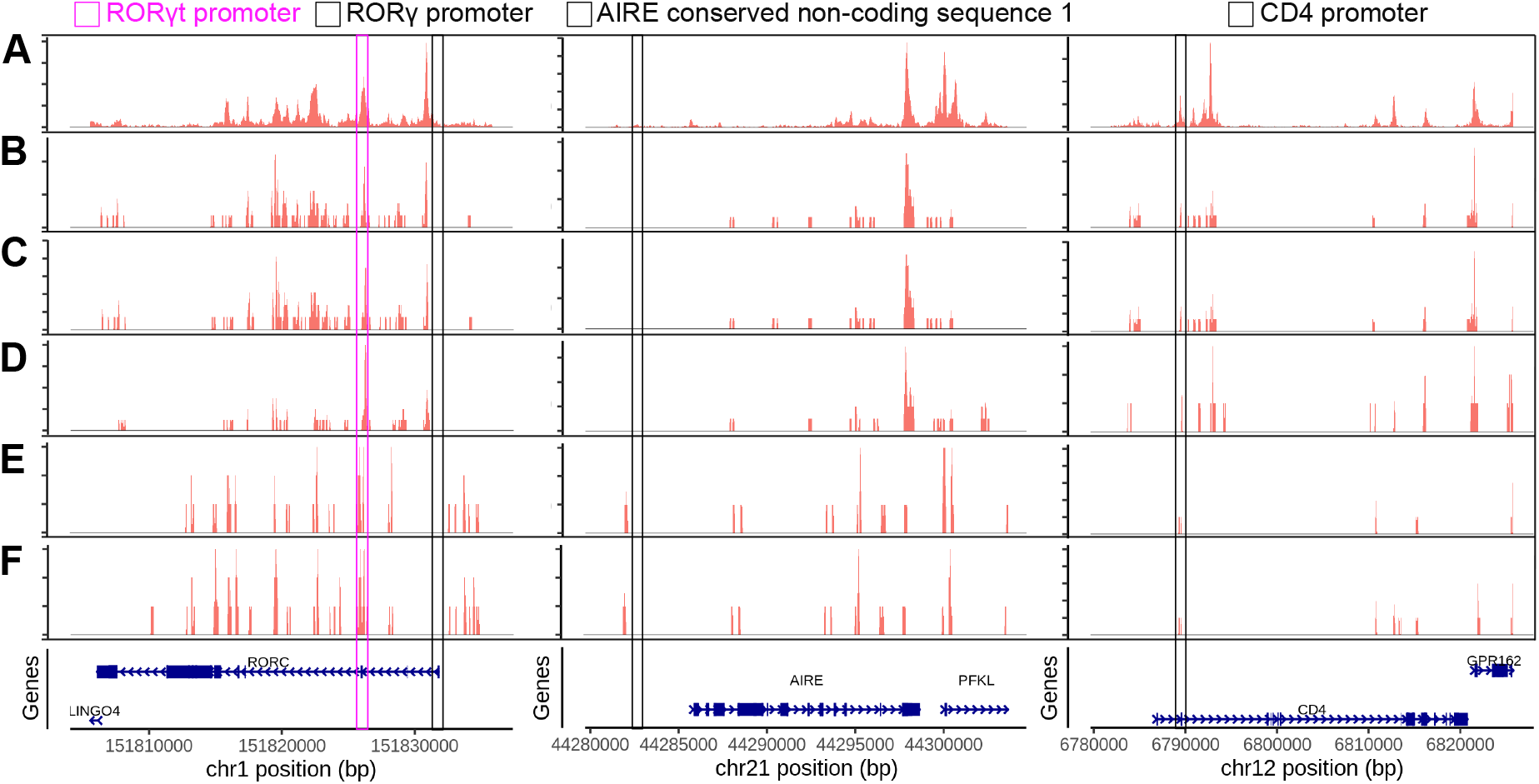
Chromatin accessibility at the *RORC, AIRE*, and *CD4* loci in R-DC-like cells. (A– F) Accessibility from the *RORC* locus (left column), *AIRE* locus (middle column), and *CD4* locus (right column) from the paired Ulezko Antonova *et al*. dataset that originally described these R-DC-like cells ^49^ (A), paired tonsil cell atlas data ^51^ (B), scPairing’s re-pairing of the paired tonsil cell atlas (C), scPairing’s artificial pairing of the unpaired tonsil cell atlas (D), paired intestine data ^52^ (E), and scPairing’s re-pairing of the paired intestine data (F). In the *RORC* column, the pink box indicates the *RORγt* promoter region and the black box indicates the *RORγ* promoter. In the *AIRE* column, the black box indicates a conserved non-coding sequence of *AIRE*. In the *CD4* column, the black box indicates a *CD4* promoter region.

We first artificially paired data from an atlas of human tonsil cells.^51^ This data contains both paired and unpaired scRNA-seq and scATAC-seq cells. Starting with the paired data, we verified that scPairing correctly re-pairs the R-DC-like cells. Since the two modalities were already paired, we checked whether each scRNA-seq profile of an R-DC-like cell paired with an scATAC-seq profile of an R-DC-like cell. Out of 27 R-DC-like scRNA-seq profiles, 24 paired with a scATAC-seq profile from a R-DC-like cell, including 12 perfect matches (**Supplementary Table S4**). Comparing the chromatin accessibility profiles, we found that the profiles of our re-pairings matched the profiles from the original paired data and the Ulezko Antonova *et al*. data at all four key regions (**Fig. 5B and 5C**).

Given the success of scPairing at re-pairing rare R-DC-like cells, we then applied scPairing to the unpaired cells. Examining the chromatin accessibility profiles paired to R-DC-like scRNA-seq profiles, we see that they mirror both the Ulezko Antonova *et al*. dataset and the original paired tonsil atlas dataset at the four regions of interest (**Fig. 5D**).

Next, we applied scPairing to paired human intestine cells^52^ to show that our pairing procedure works across tissues. The chromatin accessibility profiles of the original paired data and our re-pairings were consistent with each other (**Fig. 5E and 5F, Supplementary Table S5**). The *RORC* and *CD4* loci were consistent with the Ulezko Antonova *et al*. dataset, but the *AIRE* CNS1 exhibited accessibility close to the region in both the paired and re-paired data. This different accessibility profile with *AIRE* CNS1 accessibility has been observed in the mouse spleen.^53^ *Aire* has also been shown to be highly expressed in R-DC-like cells in mouse mesenteric lymph nodes, which supports *Aire* CNS1 accessibility as the region contains NF-*κ*B binding sites critical for *Aire* expression.^54^,^55^scPairing’s success in pairing rare cells both within the same tissue and across tissues shows its ability to generate multiomics profiles of underrepresented cells to better understand their cellular state.

### Applying scPairing re-annotates clear cell renal cell carcinoma data and generates new multiomics profiles

One challenge with applying our pairing framework is the lack of a single paired multiomics dataset that captures all cell types in the unpaired data in sufficient numbers. To demonstrate how we alleviate this challenge, we applied scPairing to integrate and pair separate scATAC-seq and scRNA-seq cells from clear cell renal cell carcinoma (ccRCC) samples.^56^ We first selected a multiomics bridge dataset of 23,001 cells sequenced from ten ccRCC samples (**Supplementary Fig. S5A**).^57^ However, this dataset consists primarily of malignant cells, with different immune cells comprising a small proportion of cells at 11.5%, and other cell types represented by fewer than 50 cells. Thus, to increase the proportion of immune cells, we incorporated 10,970 PBMCs from 10x Genomics,^58^ resulting in a final bridge dataset of 33,971 cells (**Supplementary Fig. S5B**).

We first applied scPairing to re-annotate the scATAC-seq data by integrating the separate scATAC-seq and scRNA-seq modalities onto a common embedding space, followed by transferring the labels from the scRNA-seq onto the scATAC-seq data with a 5-nearest neighbors classifier (**Fig. 6A, Supplementary Fig. S5C**). The original annotations of the scATAC-seq data were based upon gene activities, differentially expressed peaks, and transcription factor analysis (**Supplementary Fig. S5D**)^56^, and only captured the major cell types (ten cell subsets) (**Fig. 6B**). We next compared the annotations obtained from scPairing to Seurat CCA (**Fig. 6C**).^45^ While CCA also corrects some of the original annotations, it misclassified plasma cells as B cells and labeled some malignant cells as plasma cells (**Supplementary Fig. S5E and S5F**), as supported by marker gene accessibility (**Supplementary Fig. S6**). Notably, scPairing corrected many of the misannotations reported in the original study, including misannotated monocytes as B cells, and misannotated B and plasma cells as fibroblasts (**Fig. 6B, Supplementary Fig. S6**), highlighting the effectiveness of scPairing in cross-modality cell type annotation.

**Figure 6:**
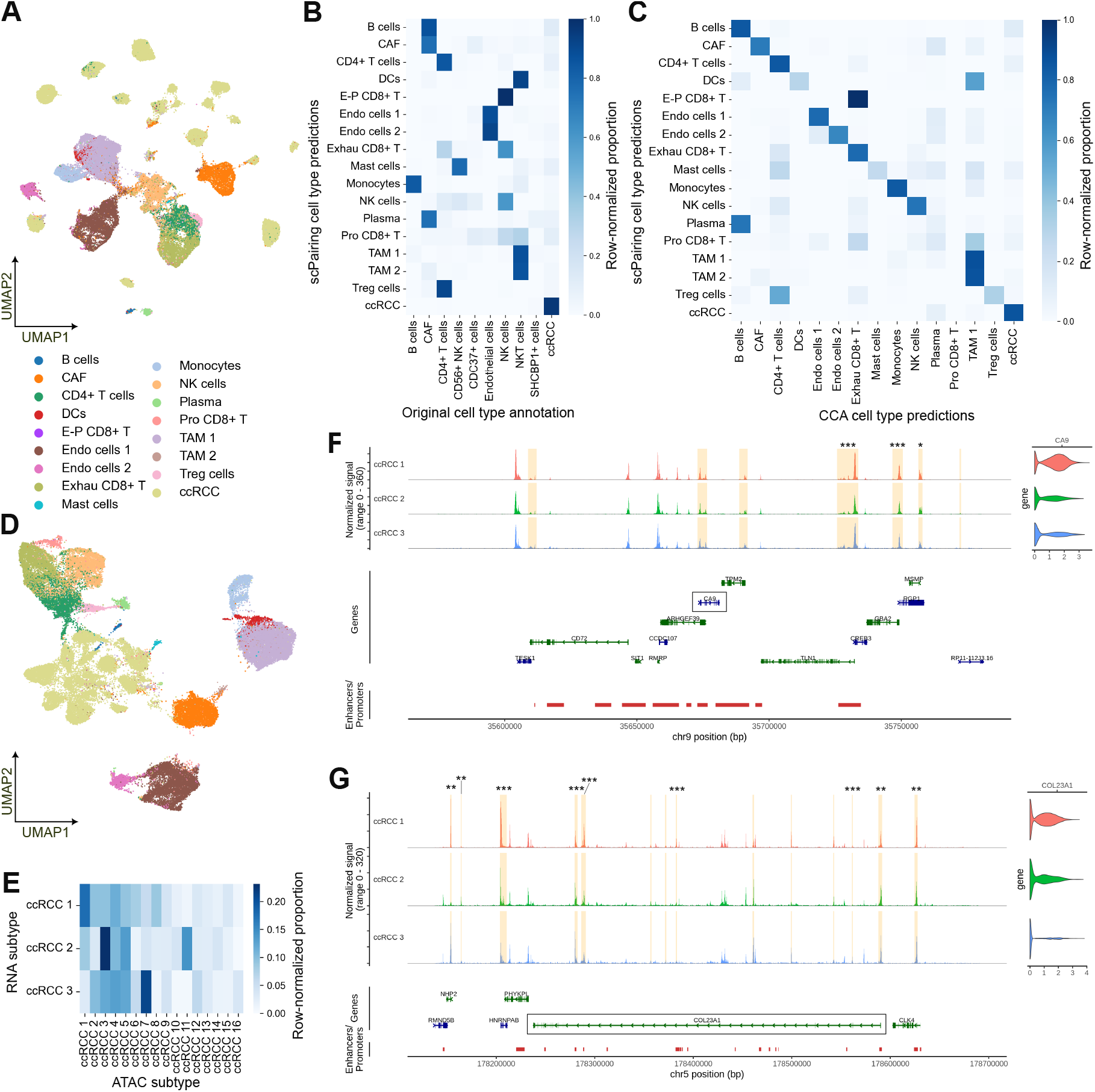
Re-annotating and generating ccRCC single-cell multiomics data with scPairing. UMAP visualization of the scATAC-seq ccRCC data from Yu *et al*., ^56^ colored by re-annotated cell types using bridge integration with scPairing. (B and C) Comparison of scPairing cell type annotations of the Yu *et al*. ccRCC scATAC-seq data ^56^ against the original annotations (B) and CCA annotations (C). The heatmaps are normalized per row. (D) UMAP visualization of the artificially paired multiomics data colored by scRNA-seq cell type annotation. (E) Heatmap of correspondences between ccRCC scRNA-seq subtypes and scATAC-seq subtypes in the artificially paired data, normalized per row. (F and G) Accessibility around the *CA9* locus (F) and *COL23A1* locus (G). The gene models for *CA9* and *COL23A1* are highlighted in a black box. Gene regulatory elements identified in GeneHancer ^60^ are displayed in red. Regions highlighted in yellow have accessibility correlated with gene expression. Highlighted regions that are differentially expressed in a ccRCC subtype are annotated with asterisks, according to a *t*-test with adjusted *p*-value after Benjamini-Hochberg correction: *p <* 0.05 (*\**), *p <* 0.01 (*\*\**), and *p <* 0.001 (*\* * \**).

Next, we generated 56,718 artificially paired multiomics profiles with scPairing (**Fig. 6D**). Leveraging the flexibility of scPairing’s bridge integration formulation, we avoided batch correction in the low-dimensional representations used as input to scPairing to preserve inter-patient heterogeneity. Based on marker genes from Yu *et al*.,^56^ we annotated three scRNA-seq subtypes and 16 scATAC-seq subtypes within the ccRCC malignant cells. Our artificial pairings showed an association between scRNA-seq subtype 2 and scATAC-seq subtype 11, as previously reported,^56^ but also showed that scRNA-seq subtype 2 was associated with scATAC-seq subtype 3, and scRNA-seq subtype 3 with scATAC-seq subtype 7 (**Fig. 6E**), which demonstrates scPairing’s potential to find associations between disease subtypes.

With our artificially paired data, we applied the SEACells algorithm^48^ to identify relationships between key ccRCC genes and peaks. We first looked at *CA9*, a ccRCC diagnostic marker present only in tumor cells^59^ that is differentially expressed in scRNA-seq subtype 1 compared to subtypes 2 and 3 (**Fig. 6F**). We identified a number of peaks that were correlated with *CA9* expression, including three peaks that were more highly expressed in scRNA-seq subtype 1. Moreover, many of the correlated peaks corresponded to known regulatory elements from GeneHancer.^60^ We then looked at *COL23A1*, a potential prognostic factor for ccRCC,^61^ where our artificially paired data revealed many candidate regulatory regions for *COL23A1* that also exhibited differential accessibility across the three scRNA-seq subtypes (**Fig. 6G**). Overall, these results show that the artificially paired data generated by scPairing can facilitate downstream analyses and biological discovery.

### scPairing extends to trimodal sequencing data

To demonstrate that scPairing can integrate more than two data modalities, we applied our model to a TEA-seq dataset, which contains joint measurements of gene expression, chromatin accessibility, and cell epitopes.^25^

The representations learned by scPairing captures major cell types and recapitulates known benefits of detecting cell epitopes, such as CD56^bright^ and CD56^dim^ natural killer cell subtypes (**Fig. 7A**).^4^ The representations for each individual modality also showed separation of cell types (**Fig. 7B**) while mixing the three modalities in the common embedding space (**Fig. 7C**). Compared to CLUE and Cobolt, scPairing embeds each modality closer to its true match across all pairs of the three modalities (**Fig. 7D**). To investigate whether scPairing can separate cells with shared profiles in one or two modalities, we shuffled two cell type’s profiles to generate eight combinations of scATAC-seq, scRNA-seq, and epitope data (**Supplementary Fig. S7A**). After applying scPairing, we observed that all eight combinations were separated (**Supplementary Fig. S7B–E**), suggesting that scPairing can distinguish cells with shared expression profiles.

**Figure 7:**
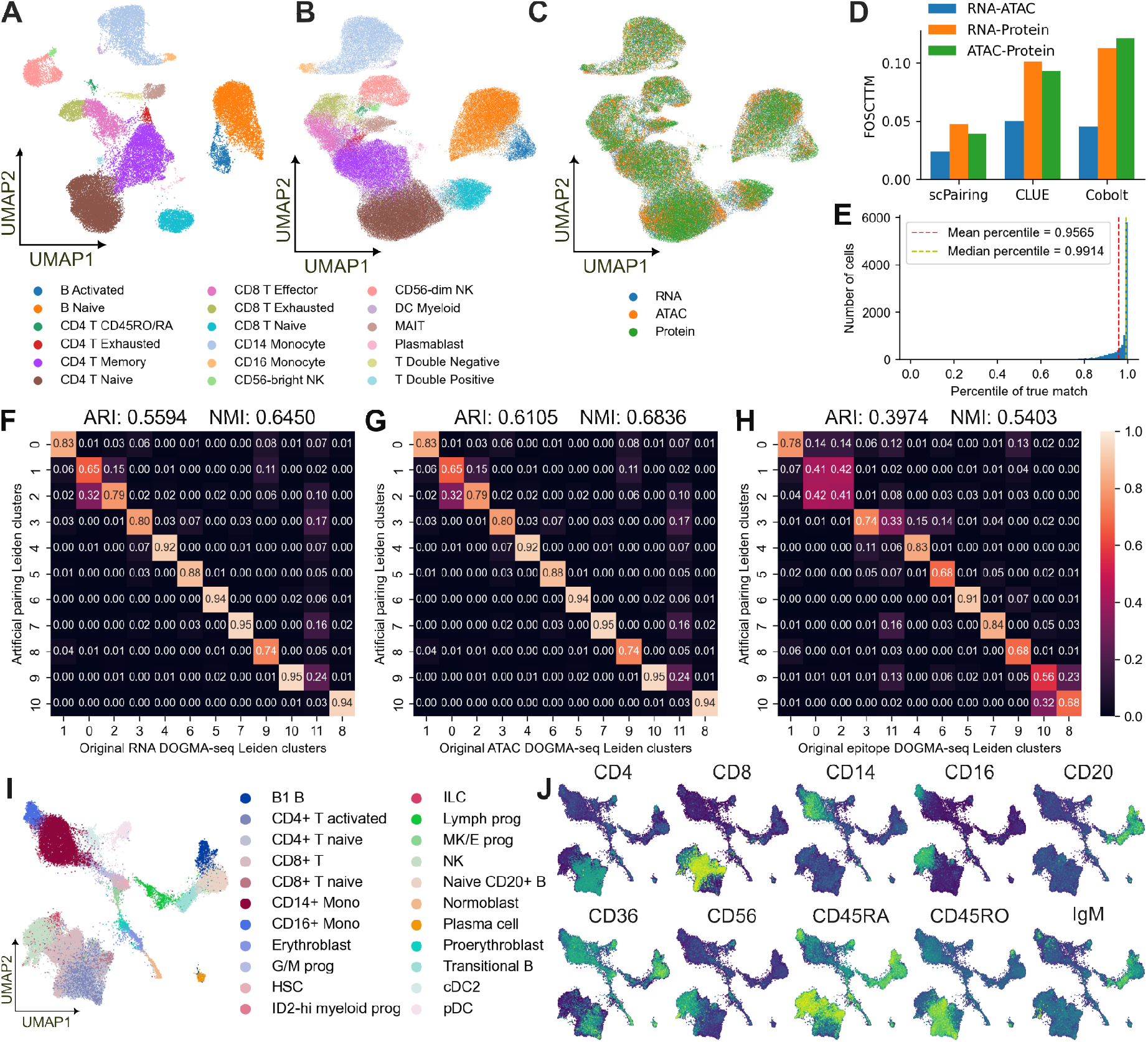
Integration of joint scRNA-seq, scATAC-seq and cell epitope data. (A) UMAP visualization of scPairing joint representations of the TEA-seq data, colored by manually annotated cell types. Each dot represents a cell. (B and C) UMAP visualization of scPairing representations of each of the three modalities in the embedding space, colored by manual cell type annotation (B) and modality (C). Each dot represents a profile from one specific modality of a cell. (D) Comparison of pairwise-modality FOSCTTM on TEA-seq embeddings. (E) Histogram of percentile rank of true match among all possible pairings for DOGMA-seq cells projected onto the TEA-seq bridge data. The yellow dashed line denotes the median percentile rank, while the red dashed line denotes the mean percentile rank. (F–H) Proportion of cells from each original DOGMA-seq Leiden cluster in each re-pairing Leiden cluster. The matrix is normalized by column; each column sums to one. Since the re-pairings constitute three different sets of cells, one for each modality, we computed the similarity for the selected scRNA-seq cells (F), scATAC-seq cells (G), and epitope cells (H). (I) UMAP visualization of the scRNA-seq modality of the artificially generated trimodal BMMCs, colored by cell type. (J) Marker epitope expression in the artificially paired trimodal BMMCs.

Next, we applied scPairing to re-pair DOGMA-seq data^24^ using the TEA-seq data as a bridge. The DOGMA-seq embeddings produced by scPairing well preserve the matching triplets, as the true matching ranks very highly in terms of embedding similarity among all possible matchings (**Fig. 7E**). The true matching triplets also exhibited high pairwise cosine similarity and low FOSCTTM, indicating that they co-embed closely (**Supplementary Table S6**). The scRNA-seq and scATAC-seq modality pair showed better alignment metrics, suggesting that finding the true pairing with epitope data is harder. This difficulty may stem from fewer overlapping features between datasets, with only 39 proteins shared by the TEA-seq and DOGMA-seq data, while both the scRNA-seq and scATAC-seq modalities have much greater overlap.

Then, we used a randomized greedy algorithm to generate a re-pairing of the DOGMA-seq data, as no efficient optimal algorithm exists for this problem (**Methods**). Next, we evaluated whether our re-pairing reflects the original data. Without ground-truth cell type labels, we compared the similarity of clusters discovered in the original data versus the repaired data. The cells in the re-pairing contain three modalities combined to form one cell, each of which may originate from different cells. Thus, we computed the similarity with respect to each modality. We observed that the clusterings show strong similarities, with substantial overlap regardless of modality (**Fig. 7F–H**), with an average adjusted Rand index (ARI) of 0.522 and normalized mutual information (NMI) of 0.623.

Finally, we synthesized a new trimodal dataset using the joint scRNA-seq and scATAC-seq Multiome BMMC benchmarking dataset and the epitopes from the CITE-seq dataset from Luecken *et al*.,^38^ using the TEA-seq data as a bridge. After projecting all three modalities onto the bridge, we assigned epitope profiles to the Multiome cells. Out of 69,249 Multiome cells, we successfully paired 56,569 cells to high-similarity epitope profiles (**Fig. 7F**). Despite the TEA-seq bridge missing cell types present in the BMMC data, we still found pairings for the unobserved erythroid cell types, which suggests that scPairing could learn an embedding space that matches out-of-distribution cells. We also observed that the matched epitope profiles are consistent with the cell identities through the expression of marker epitopes, including the separation between CD4 and CD8 expressing cells, and CD45RA and CD45RO expressing cells (**Fig. 7G**).

## 3 Discussion

In summary, scPairing builds upon the powerful variational autoencoder architecture by adding contrastive learning to align representations of multiomics data together onto a common embedding space. By learning the alignment between paired multiomics data, scPairing enables the alignment of unpaired data, from which new artificial multiomics data can be generated by pairing cells together.

Generating artificial pairings with scPairing can be viewed as a reformulation of bridge integration. As originally defined, bridge integration expresses unimodal cells in terms of multiomics cells (the bridge) to harmonize the features for cells from separate modalities using dictionary learning.^20^ While both our method and the original bridge integration first harmonize the bridge with the unimodal data, our method differs by using the bridge data to find an alignment between the modalities. This procedure is more efficient than dictionary learning, which expresses unimodal cells in terms of bridge cells.

Unlike many other multiomics integration tools taking raw counts as input, scPairing can take low-dimensional representations of each modality as inputs. This strategy reduces scPairing’s runtime, although computing the modality-specific representations can be computationally expensive for complex deep learning models. However, the availability of pre-trained models, including pre-trained scVI models^32^ or single-cell LLMs,^33^,34,62 each trained on millions of cells, helps alleviate this issue. These pre-trained models can compute embeddings either after fine-tuning or directly in a zero-shot manner without any additional training, drastically reducing the time required to obtain input representations for scPairing. Our benchmarking with scGPT^33^ and CellPLM^34^ embeddings demonstrates that these pre-trained models can be used with scPairing. As large pre-trained models designed for chromatin accessibility become available, such as EpiGePT,^63^ we foresee scPairing interfacing directly with pre-trained models for any modality, allowing for fast multiomics data integration.

The flexibility in choosing modality-specific integration methods as input for scPairing opens the door to cross-species multiomics data generation. Fewer non-human paired multiomics data exist due to the high cost of multiomics sequencing. Given the success in pairing R-DC-like cells using bridge and unimodal data from different tissues, one can potentially also apply scPairing to bridge and unimodal data from different species. Specialized tools developed for cross-species integration can assist in integrating the bridge and unimodal data.^64–66^

We believe that scPairing presents an opportunity to leverage the wealth of unimodal single-cell data to generate high-quality artificial multiomics data, including difficult to obtain trimodal data. These generated datasets can be analyzed through a multiomics lens using other powerful multiomics tools to identify novel relationships between modalities.

### Limitations of the study

There are some challenges in applying scPairing. First, the linear sum assignment algorithm used to generate pairings in the common embedding space does not scale efficiently when pairing large unimodal data with many similar cells, such as the retina data. This limitation requires approximation by computing pairings on separate chunks of the data. Approximation algorithms for linear sum assignment could improve the scalability of pairing.^67^ Second, our pairing framework requires a multiomics dataset to use as a bridge, which may not exist within certain contexts. In these cases, methods for diagonal integration may be more appropriate as they do not require the datasets to have common features.

## 4 Methods

### The scPairing framework

Though scPairing applies to any combination of data modalities, we describe the two-modality version of scPairing in terms of joint scATAC-seq and scRNA-seq data. The scPairing model takes in a multimodal dataset with gene expression matrix 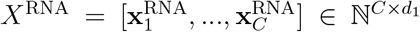, and chromatin accessibility matrix 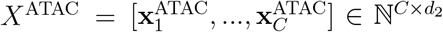, with *C* cells, *d*_1_ genes, and *d*_2_ peaks. scPairing also takes in a batch-corrected low-dimensional representation of each modality, 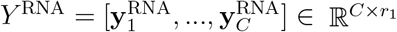 and 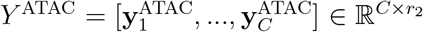, with *r*_1_ and *r*_2_ being the dimensions of the representations.

These representations can be obtained from approaches such as PCA for gene expression data and LSI for chromatin accessibility data. Alternatively, these representations can also be obtained from deep learning models such as scVI^32^ for gene expression data and PeakVI^35^ or cisTopic^68^ for chromatin accessibility data, or from pre-trained models based on transformers.^33^,^34^,^69^

In scPairing, we incorporate the Contrastive Language Image Pre-training (CLIP) model described by Radford *et al*.^36^ designed for joint embedding of image and text representations into a variational autoencoder model. For each cell i, we assume that the representations 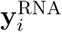 and 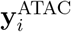 are described by a low-dimensional latent vector **z**_*i*_ ∈ ℝ^*d*^ that lies on a (*d* − 1)-sphere *S*^*d*−1^,

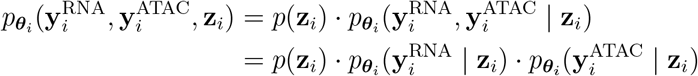

where ***θ***_*i*_ is the parameter set for each distribution. The prior distribution *p*(**z**_*i*_) is assumed to be the uniform distribution on the (*d*− 1)-sphere. Both 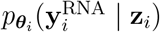 and 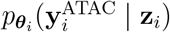 are assumed to be normally distributed, with parameters specified by neural networks (decoders).

We use variational inference to approximate the posterior distribution for **z**_*i*_. The variational distribution 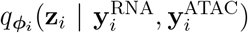 is assumed to be a Power Spherical distribution^70^ on a (*d*− 1)-sphere. The Power Spherical distribution is an alternative to the von Mises-Fisher (vMF) distribution, which is considered the normal distribution on the hypersphere. We chose the Power Spherical distribution because of its sampling efficiency compared to the vMF distribution. The Power Spherical distribution has two parameters, the direction ***µ*** ∈ *S*^*d*−1^ and the concentration *κ* ∈ ℝ_*>*0_.

This variational distribution is parameterized by the three neural network encoders *f*_RNA_, *f*_ATAC_ and *f*_*κ*_, which compute ***µ***_RNA_, ***µ***_ATAC_, and *κ*, respectively. Since *f*_RNA_ and *f*_ATAC_ both produce a *d*-dimensional direction, we merge the directions from both encoders using a method derived from dropout^71^,^72^to produce the ***µ*** for the Power Spherical distribution. For each index *i* ∈ {1, 2, *· · ·, d}*, we take the value corresponding to the *i*-th index in ***µ***_RNA_ with 0.5 probability. Otherwise, we take the value corresponding to the *i*-th index in ***µ***_ATAC_.

This approach allows the model to better generalize to varying emphases on ***µ***_RNA_ or ***µ***_ATAC_. The evidence lower bound (ELBO) objective for training both encoder and decoder neural networks becomes

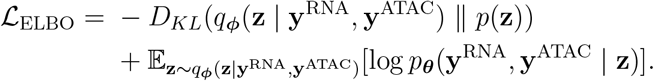

Here, 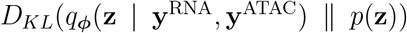 represents the Kullback-Leibler (KL) divergence between 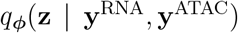 and *p*(**z**). We also remove the cell index *i* here for brevity, assuming just one given cell. Given *N* cells, the final ELBO is the sum of the individual ELBO over *N* cells.

Given a Power Spherical distribution *P* and a uniform distribution on the sphere *Q* = **U** (*S*^*d*−1^), the KL divergence has a closed form solution,^70^

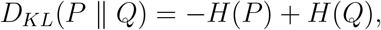

where *H* is the entropy of the distribution. Since we assumed the prior distribution *p*(**z**_*i*_) to be a uniform distribution on the hypersphere, we can compute the closed-form solution for the KL divergence between the variational approximation and the prior distribution.

To enforce that the two embeddings of a cell are close on the hypersphere, the encoders are learned to minimize the CLIP contrastive loss. In each batch of *N* cells, scPairing maximizes the cosine similarity of the embeddings for the *N* matching measurements, while minimizing the cosine similarity of the embeddings for the *N* ^2^ − *N* non-matching measurements. For each RNA direction 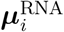, we compute the cross-entropy loss and average over all cells,

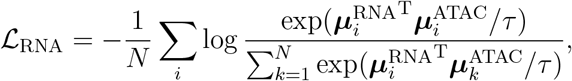

where *τ* is the learned temperature parameter. For each ATAC direction 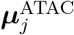, we compute the symmetric version of the previous loss and average over all cells,

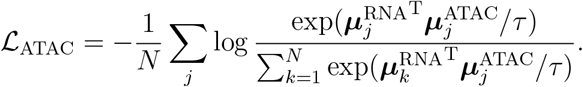

In addition to the contrastive objective, we add an adversarial objective by including a discriminator neural network *h* : *S*^*d*−1^ *→* [0, 1] that takes each direction, 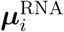 or 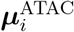, as input and outputs the probability that it originates from an RNA measurement. The RNA and ATAC encoders are trained to fool the discriminator such that the discriminator cannot determine the difference between the RNA embeddings and ATAC embeddings. The adversarial loss is given by

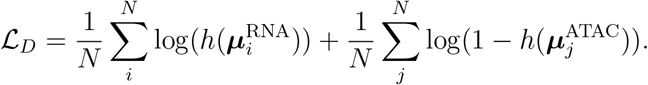

Finally, we add an alignment objective to further encourage modalities from the same cell to be similar,

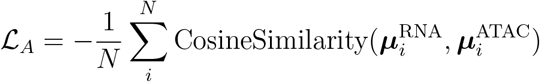

In datasets where batch effects are prominent and cannot be sufficiently corrected by the modality-specific encoders, the CLIP contrastive loss may over-separate cells based on their batch. To counteract this effect, we add another adversarial objective, except this discriminator predicts the batch each embedding came from. Given a dataset with *B* batches, the discriminator neural network *b* : *S*^*d*−1^ *→* [0, 1]^*B*^ takes in 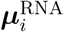 or 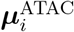 and produces the probabilities of cell *i* coming from each batch. The RNA and ATAC encoders are trained to fool this discriminator. The batch adversarial loss is a *B*-class cross-entropy loss,

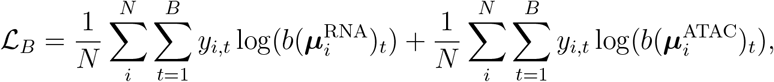

where *y*_*i,t*_ is a binary indicator for cell *i* belonging to batch *t*.

The overall objective of scPairing is:

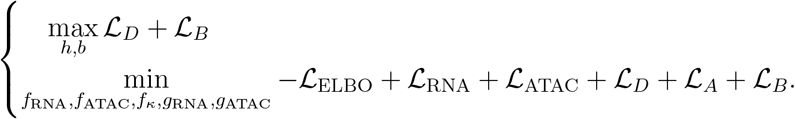

### Extending scPairing to allow for full data reconstruction

While the model described so far works for producing aligned embeddings usable in downstream analyses, the model does not generate raw count data, unlike other methods.^17^,^18^,^22^To enable data imputation, we augment the model with two additional decoders that take the reconstructed representations 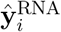 and 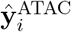 and reconstruct the count data 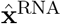 and 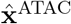. We also define probabilistic models of scRNA-seq and scATAC-seq count data to modify the scPairing ELBO.

We model the scRNA-seq counts with a negative binomial distribution. For each observed count 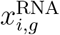 of gene *g* in cell *i* the scRNA-seq data matrix 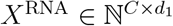, we model 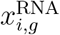 as

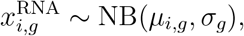

where the mean *µ*_*i,g*_ is specified by a neural network decoder *ĝ*_RNA_, while the dispersion parameter *σ*_*g*_ is a gene-specific non-negative number shared across cells.

We model the scATAC-seq counts with a Bernoulli distribution. For each observed count 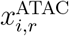 of region *r* in cell *i*, we binarize the observation into either zero or one. This binarized count is 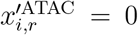 if 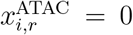 and 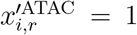 if 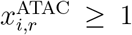. Then, we model this observation as

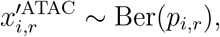

where the probability parameter *p*_*i,r*_ is specified by a different neural network decoder *ĝ*_ATAC_.

The new ELBO formulation becomes

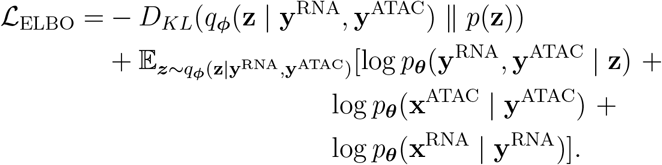

### Leveraging pre-trained decoders for full data reconstruction

Instead of using a decoder that is trained from scratch during the training process, if the low-dimensional representations *Y* ^RNA^ or *Y* ^ATAC^ were computed using a method that allows for reconstruction, we can leverage those decoders for faster training. Methods such as PCA and variational autoencoders already contain a reconstructor, either by multiplying the representations by the principal axes in PCA, or passing the latent representations through the VAE decoder. Instead of learning new decoders to reconstruct the counts, we can pass *Ŷ*^RNA^ and *Ŷ*^ATAC^ into the reconstructor from the method originally used to compute *Y* ^RNA^ or *Y* ^ATAC^.

### Extension to datasets with more than two modalities

To extend scPairing to *M* modalities with count matrices *X*^(1)^, *X*^(2)^, …, *X*^(*M*)^ and batch-corrected low-dimensional representations *Y* ^(1)^, *Y* ^(2)^, …, *Y* ^(*M*)^, we add additional encoders and decoders for the extra modalities, and compute the contrastive loss between all pairs of modalities. There are encoders *f*_1_, …, *f*_*M*_, where *f*_*m*_ takes in *Y* ^(*m*)^ as input and produces *d*-dimension direction ***µ***^(*m*)^, and decoders *g*_1_, …, *g*_*M*_, where *g*_*m*_ takes in the merged *d*-dimension embedding and reconstructs *X*^(*m*)^. The process to merge the *M d*-dimension directions from each encoder uses the same dropout method, except each encoding has a 1*/M* chance of being selected at each index. The pairwise CLIP loss between modality *m*_1_ and *m*_2_ is given by

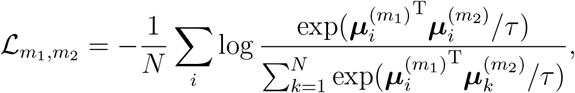

where this loss is calculated across all modalities *m*_1_, *m*_2_, where *m*_1_ ≠ *m*_2_. The discriminator *h* changes from a function *h* : ℝ^*d*^ *→* [0, 1] to *h* : ℝ^*d*^ *→* [0, 1]^*M*^, a function that outputs the probabilities that the embedding came from the *m*-th modality for *m* ∈ {1, …, *M*}. The discriminative loss becomes an *M* -class cross-entropy loss,

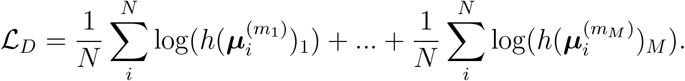

The alignment objective becomes a pairwise cosine similarity between all modalities,

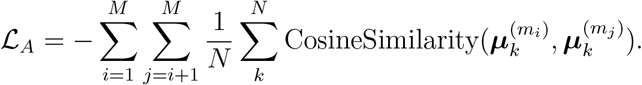

The batch discriminator objective is modified to calculate the prediction across all modality embeddings,

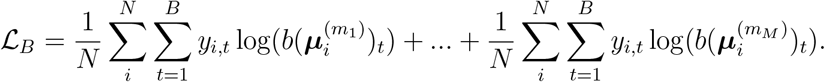

The new objective becomes:

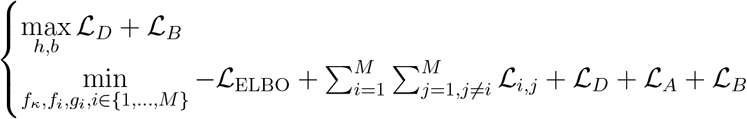

### Model structure and training

All of the neural network components of our model use a single-layer neural network with 128 units, layer normalization, and exponential linear unit activation.^73^ Training is a two-step process due to the two objectives, training the discriminators to maximize the adversarial objectives, and then training the encoders and decoders to minimize the augmented VAE objective. We use the Adam optimizer^74^ during training, with an initial learning rate of 0.005 and learning rate decay of 0.00006. We use a minibatch size of either 2000 or 5000 since larger minibatch sizes are better for representation learning tasks.^75^

### Artificially pairing unpaired multiomics data

Given a joint scRNA-seq and scATAC-seq bridge dataset *R* with matrices 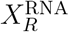 and 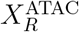, scPairing can generate pairings between unpaired scRNA-seq dataset *P* and scATAC-seq dataset *Q* with matrices 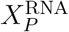 and 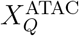. As introduced by Hao *et al*.,^20^ we first compute a common batch-corrected low-dimensional representation for each modality across datasets. For the scRNA-seq data, 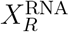 and 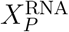 are processed into 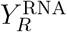 and 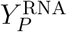, where batch effects between *R* and *P* are resolved. Likewise, 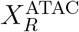 and 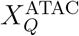 are processed into batch-corrected representations 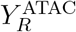 and 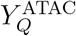. Then, we train scPairing on the bridge dataset 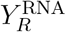 and 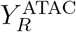. With trained RNA and ATAC encoders, we pass 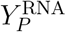 through the RNA encoder and 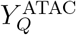 through the ATAC encoder to obtain embeddings 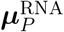 and 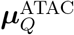 on the common hyper-spherical space.

The problem of pairing these cells is treated as a maximum weight matching problem in bipartite graphs, which aims to find a pairing between vertices from one partition to the other such that the sum of edge weights is maximized. We first generate a bipartite graph between cells *i* from dataset *P* and *j* from dataset *Q*, with the weight of each edge between cells *i* and *j* being

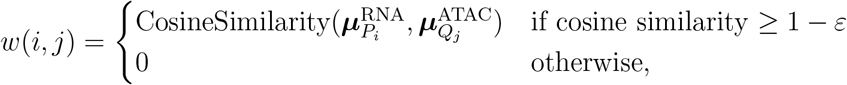

for some cutoff adjustment *ε >* 0. We prune edges between cells with similarity less than 1−*ε* to prevent dissimilar cells from being paired together during matching. The *ε* parameter can be manually tuned in the context of the data. In data with many similar cell types, choosing a smaller *ε* will prevent similar but distinct cell states from pairing together, such as cells present along a differentiation trajectory.

To solve the maximum weight matching problem, we use the linear sum assignment function from SciPy,^76^ which uses a modification of the Jonker-Volgenant algorithm.^77^ In large datasets, computing the maximum weight matching is infeasible due to time and memory constraints. In such cases, we find an approximate solution by dividing the cells in both datasets into random disjoint subsets and computing the maximum weight matching on each subset.

### Trimodal artificial pairing

Given a joint scRNA-seq, scATAC-seq and epitope dataset bridge dataset *R* with matrices 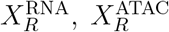, and 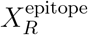, and corresponding unpaired data from datasets *P, Q*, and *S* with matrices 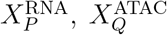, and 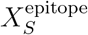, we can apply scPairing to embed all three modalities onto the common hyperspherical space using the trimodal extension of scPairing. This procedure follows from pairing two modalities, except a tripartite graph is produced, with one partition each for the scRNA-seq, scATAC-seq, and epitope modality cells.

The problem of finding artificial pairings for three modalities is intractable as finding a maximum tripartite graph is NP-hard. In lieu of finding an optimal pairing, we propose a randomized greedy algorithm to generate a respectable pairing. Our algorithm randomly selects a cell *i* from the modality with the fewest cells (without loss of generality, assume the scRNA-seq dataset has the fewest cells) and finds the cell *j* from the scATAC-seq dataset and cell *k* from the epitope dataset that maximizes the pairwise cosine similarity between 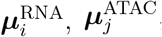, and 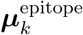. Cells *i, j*, and *k* are removed from consideration and the algorithm repeats until all cells are paired or the pairwise cosine similarity drops below a threshold. The full algorithm pseudocode is provided in **Supplementary Note S1**.

Alternatively, if we have a joint scRNA-seq and scATAC-seq dataset *P* with a separate epitope dataset *Q*, we can also generate trimodal data. In this case, all three modalities are still embedded on the common hyperspherical space learned by scPairing with bridge dataset *R*, except we can make a bipartite graph with one partition being the paired scRNA-seq and scATAC-seq cells, and the other partition being the epitope cells. The edge weights between cell *i* and cell *j* is

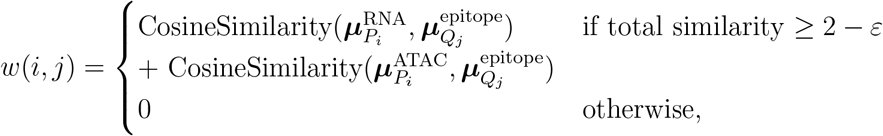

for some cutoff adjustment *ε*. With this formulation, we can find optimal pairings using the linear sum assignment algorithm.^76^ This pairing method can also be applied if we have a joint scRNA-seq and epitope dataset with a separate scATAC-seq dataset.

### Benchmarking evaluation

We assessed multiomics data integration using different metrics for data integration and alignment on the NeurIPS 2021 dataset.^38^ The evaluation metrics are described below.

### Conservation of biological structure using kNN accuracies

To compare the quality of embeddings produced by each model, we performed both cross-batch validation and tenfold cross-validation of cell type prediction. Cross-batch cell type prediction takes batch correction into account, as batches must be integrated together for accurate classification. Conversely, ten-fold validation does not account for batch effects. We conducted the *k*nearest neighbor predictions with *k* = 5, 17, 29, 41, 53, and 65. For the human PBMC CITE-seq dataset, we used the second-level annotations.

### Cell matching

To evaluate whether the model can correctly embed matching measurements from the same cell, we calculated the FOSCTTM.^40^ The FOSCTTM metric indicates the proportion of cells that are closer to the embedding of one modality of a cell than the embedding of the other modality for that same cell. Since FOSCTTM is not symmetric, the final metric is the average between the two FOSCTTMs. A lower FOSCTTM indicates better cell matching.

### Modality mixing

We check that the embeddings for each modality in the multiomics dataset mixes together using the Local Inverse Simpson’s Index (LISI) metric. We use the graph iLISI implementation from scIB.^37^

### Cell type classification

We evaluate whether the representations of the unimodal data during bridge integration can enable cell type classification from the scRNA-seq data onto the scATAC-seq data. We split the benchmarking data into the paired bridge, consisting of three out of four sequencing sites, and use the separate scRNA-seq and scATAC-seq data from the fourth site as the unimodal data. Each method is trained on the bridge data and the unimodal data is directly passed into the trained model to obtain separate scRNA-seq and scATAC-seq embeddings. We use a 5-nearest neighbor classifier trained on the unimodal scRNA-seq data embeddings and predict the cell type using the scATAC-seq data embeddings. We assess the classification accuracy.

### Cross-modality imputation

We evaluate the ability of each model to impute across modalities by training the model on three out of four sites and imputing using the held-out site. We quantified the scRNA-seq imputations using the Spearman correlation between the true and imputed expression. We quantified the scATAC-seq imputations using the binary cross-entropy between the binarized true accessibility and the imputed accessibility.

### Comparison methods

We compare our method to nine other multiomics tools: GLUE,^21^ MOFA+,^19^ MultiVI,^17^ MoETM,^15^ MIRA,^12^ Cobolt,^18^ CLUE,^22^ BABEL,^29^ and scButterfly.^47^ Of these, Cobolt, CLUE, and GLUE were designed for modality alignment, and BABEL and scButterfly were designed for inter-modality imputation. We ran the methods based on their publicly available tutorials, or if no tutorials exist, based on their default settings. All tools were installed from publicly available source code, with the exception of BABEL, which was re-implemented to interface with AnnData.

For our experiments benchmarking GLUE using the CITE-seq data, we manually construct a graph, termed the prior regulatory network. We annotate the Ensembl identifiers of the genes responsible for a given protein to add protein-gene edges. In the case of protein complexes comprised of multiple genes, we connect all of them to the same protein. Further, we add nodes for all genes and proteins in the dataset, as well as self-loops, to satisfy GLUE’s graph checking criteria. The weight and sign of all edges were set to 1.

### Evaluating the quality of retina pairings

With unpaired data, no ground truth pairing exists to evaluate the quality of the computed pairings. One common analysis performed on joint chromatin accessibility and gene expression data is inferring transcription factor binding by identifying correlated genes and peaks. Thus, we compare the preservation of gene-peak correlations found in a true multiomics dataset to indicate realistic pairings.

We compare our pairings against four other methods. The first is gene activities computed with Signac,^46^ which directly transforms scATAC-seq fragments into gene counts by counting fragments overlapping each gene plus 2kb upstream. The second method uses canonical correlation analysis implemented in the Seurat^45^ `FindTransferAnchors` function to integrate the gene activities onto the scRNA-seq data. Then, we use `TransferData` to impute the gene expression of the scATAC-seq cells using the scRNA-seq data as a reference. This second method is the strategy applied by Liang *et al*. in their analysis of gene-peak correlations. The last two methods are both deep learning methods that impute expression by decoding from a shared latent space: BABEL^29^ and scButterfly.^47^ We ran all of these methods on the same set of scATAC-seq cells that were present in our artificial pairings.

We focus on the retinal bipolar cells as they were also analyzed for gene-peak correlations in Liang *et al*.^43^ *In particular, Liang et al*. analyzed the correlation of gene *GNB3* expression and peak chr12:6959945-6962406 accessibility across all retina bipolar cell types. We compare the means and standard deviations of gene-peak correlations in twelve retina bipolar cell subtypes. We choose to compute correlations for all genes with peaks within 250kb of the gene TSS. Since not all genes are imputed by gene activities, we only compute gene-peak correlations for genes present in the multiomics data, the artificially paired data, and gene activities.

We use metacells, specifically the SEACells algorithm, to compute gene-peak correlations as it has been shown that metacells more faithfully capture relationships between the two modalities.^48^ We opt against donor-aggregated metacells as performed in Liang *et al*. because donor-aggregation makes the assumption that the population of cells from each donor represents a different state compared to the other donors. While this assumption may hold for data with high inter-donor heterogeneity, other data have relatively small differences between donors. In this data, scATAC-seq profiles show relatively high mixing between donors and for most cell subtypes, the bulk of cells have a diverse set of neighboring donors in the LSI embedding space, with only the FMB and IMB subtypes showing low donor mixing. Thus, when aggregating cells by donor, the difference between each metacell is likely noise, which implies that the positive, neutral, or negative correlations seen are derived through randomness between donors, not biological variation. Altogether, these observations support our choice to use SEACells aggregation.

To obtain a sufficient number of metacells to compute meaningful correlations, we run SEACells using a target of ten cells per metacell. We modify the maximum iterations of SEACells to 400 to ensure convergence, but use otherwise default parameters.

We quantify the difference between the gene-peak means and standard deviation distributions using optimal transport, which measures the distance between two distributions. We calculate both the Earth Mover distance and Sinkhorn distance using the Python Optimal Transport library.^78^ As the optimal transport is not scalable to the number of gene-peak pairs, we randomly sample 2% of the gene-peak pairs to perform optimal transport, and repeat for 100 trials.

### Evaluating scPairing robustness

In the first two robustness experiments, we used the benchmarking BMMC dataset. The ablated training dataset excluded the data from the first site, which was instead reserved as the test dataset. We then applied the bridge integration procedure using the ablated training dataset as the bridge and the test dataset for pairing.

In the first robustness experiment, we downsampled each cell type by reducing a cell type’s proportion to 2%, 1%, 0.1%, or 0% of the bridge dataset. If a cell type’s original proportion was already below 2%, we skipped that downsampling level for that cell type. We conducted two evaluations. The first evaluation uses the FOSCTTM metric from the cell matching evaluation. The second evaluation computes the proportion of cell type-consistent pairings of the downsampled cells when re-pairing the test data. A cell type-consistent pairing is one whose scRNA-seq profile and scATAC-seq profile both originated from cells of the same cell type. We compute the number of cell type-consistent pairings in the re-paired test data divided by the original number of the downsampled cell type in the test data.

In the second robustness experiment, we selected one cell type in the bridge dataset to be dominant and downsampled all other cell types such that the selected cell type composed 70%, 80%, 90%, or 95% of the total bridge cells. We repeated this procedure for the 11 most abundant cell types in the bridge dataset and evaluated the FOSCTTM in the test data separately for the dominant cell types and for the remaining cell types.

In the third robustness experiment, we evaluated the necessity of the three additional loss components by running scPairing with each combination of the three losses. In the pairing yield experiments, we used *ε* = 0.05 for the BMMC test dataset and *ε* = 0.02 for the retina data.

### Trimodal pairing evaluation

We evaluate the similarity of the re-pairings to the original pairings using three methods: FOSCTTM, the percentile rank of the true pairing among all possible pairings, and the similarity between clusters in the original data and re-paired data. The FOSCTTM is calculated as the average FOSCTTM of the three pairs of unique modalities.

The percentile rank of the true pairing calculates the cosine similarity rank of the true pairing, scRNA-seq, scATAC-seq, and epitope data all from cell *i*, among all possible pairings. Let 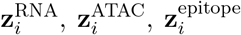, be the embeddings for cell *i* for each of the three modalities. We define the triplet similarity for cells *i, j*, and *k* to be

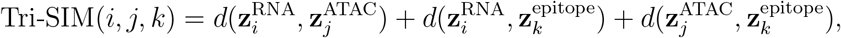

where *d* is a similarity measure (in this case, cosine similarity), as the embeddings are located on the unit hypersphere. Then, for cell *i*, we calculate the percentile rank of Tri-SIM(*i, i, i*) in

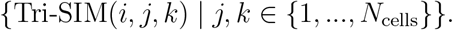

A high percentile rank triplet indicates that the true triplet match is very close to the best possible match, meaning the embeddings closely follow the true pairings.

Cluster similarity is measured using the ARI and NMI metrics. High ARI and NMI indicate that clusters formed in the re-pairings and pairings share commonalities, and the structure of the original data is preserved in the re-pairings. The ARI and NMI are calculated for each modality in the re-pairing as one artificially paired cell is composed of three cells, one for each modality.

### Data preprocessing and pairing

We detail the preprocessing steps we took for each dataset used and the parameters used for pairing.

### NeurIPS 2021 BMMC benchmarking dataset

This dataset contained 69,249 joint scRNA-seq and scATAC-seq cells and 90,261 CITE-seq cells from ten donors across four sites.^38^ Due to repeat samples from donors at different sites, there are 13 batches present in the data.

For the scRNA-seq data, we computed five low-dimensional representations: 20D Harmony-corrected PCA,^39^ 50D scVI,^32^ 10D scPhere,^31^ 512D scGPT,^33^ and 512D CellPLM.^34^ The scGPT representation was computed following their reference mapping tutorial using the continual pre-trained model checkpoint for zero-shot cell embedding. The CellPLM representation was computed following their cell embedding tutorial using checkpoint 20230926 85M. For the scATAC-seq data, we computed two representations: 20D Harmony-corrected LSI and 50D PeakVI.^35^ The LSI embedding was computed with peaks expressed in at least 1% of cells and TF-IDF transformed counts. The PeakVI embedding was computed with peaks expressed in at least 1% of cells. For the epitope data, we computed a 50D PCA embedding.

### SHARE-seq mouse skin dataset

This dataset contained 28,429 cells sequenced by SHARE-seq.^2^ We used a pre-processed and re-annotated version by Lynch *et al*.^12^ We used PCA and LSI embeddings as scRNA-seq and scATAC-seq inputs to scPairing.

### Human cerebral cortex dataset

This dataset contained 45,549 joint scRNA-seq and scATAC-seq cells.^42^ Following the original analysis, we computed 30D PCA and 10D LSI embeddings for the scRNA-seq and scATAC-seq modalities. We excluded the first LSI dimension.

### Human PBMC CITE-seq dataset

This dataset contained 161,764 cells profiled by CITE-seq.^14^ We used both 50D Harmony-corrected PCA and 50D scVI embeddings for scRNA-seq modality and 50D Harmony-corrected PCA embeddings for the epitope modality.

### Human retina datasets

We used two human retina datasets. The first dataset is over 45,835 joint scRNA-seq and scATAC-seq cells, including both left and right retinas from four donors.^44^ The second dataset contained over 250,000 cells sequenced using snRNA-seq and 137,000 cells sequenced using snATAC-seq.^43^ The snRNA-seq samples came from six donors, of which we only use the four donors with the majority of cells (211,757 cells). The snATAC-seq samples came from 21 donors, which we all use.

We filtered the scRNA-seq data by keeping cells that had between 300 and 25,000 counts and at least 10 unique genes expressed. The scATAC-seq data was filtered by retaining cells that had total counts between 3,000 and 30,000 and a nucleosome signal below four. Peaks expressed in fewer than 0.5% of cells were also removed. We merged the scATAC-seq peaks using Signac^46^ by calling peaks with MACS2^79^ on each dataset individually and combined the two peak sets using reduce.

The individual modalities in the paired and unpaired datasets were integrated with scVI and PeakVI, for scRNA-seq and scATAC-seq respectively, using the batch and dataset as categorical covariates. During pairing, we used similarity cutoff *ε* = 0.02.

### R-DC-like datasets

We used three datasets containing R-DC-like cells. The bridge dataset contained 10,523 joint scRNA-seq and scATAC-seq human tonsil cells.^49^ The bridge dataset is enriched for DC1, DC2, CD34^+^, and Lin^-^HLA-DR^+^CD4^-^CD1c^-^ cells. The human tonsil dataset we used for artificial pairing consists of 501 joint scRNA-seq and scATAC-seq cells, 1,536 scATAC-seq cells, and 7,098 scRNA-seq cells after restriction to DCs and ILC3s. The human intestine dataset we used for artificial pairing consists of 280 joint scRNA-seq and scATAC-seq cells after restricting the data to DCs. In all experiments, we used scVI as the scRNA-seq embedding and Harmony-corrected LSI as the scATAC-seq embedding. During pairing, we used similarity cutoff *ε* = 0.5.

### ccRCC datasets

We used three datasets: one PBMC dataset and two ccRCC datasets. The bridge 10x PBMCs^58^ contained 10,970 cells and no additional pre-processing was performed. The bridge ccRCC dataset contained 23,001 cells from ten donors.^57^ The scRNA-seq profiles were already pre-processed, with profiles having fewer than 1000 UMIs or more than 10% mitochondrial counts filtered out. scATAC-seq profiles were filtered to keep profiles with between 100 and 10,000,000 counts. The cells were retained if both profiles from both modalities were retained.

The unimodal pairing dataset contained 102,723 scRNA-seq profiles and 60,279 scATAC-seq profiles, obtained from 19 donors.^56^ We removed cells with fewer than 500 genes or more than twice the median of detected genes. Cells with more than 10% mitochondrial counts were filtered out. For the scATAC-seq profiles, we kept cells with between 1000 and 20,000 peak counts, a blacklist ratio at most 0.05, a nucleosome signal at most 4, and a TSS enrichment score at most 1.

We integrated the individual modalities in the paired and unpaired datasets using scVI and PeakVI, using only the source dataset as the categorical covariate. For scVI, we only used highly variable genes. For PeakVI, we used only peaks expressed in 1% of cells. For pairing the data, we used *ε* = 0.1.

### Trimodal data

We used two trimodal datasets, TEA-seq^25^ and DOGMA-seq,^24^ with both sequencing PBMCs. The TEA-seq data contained 37,124 cells sequenced in five batches. The DOGMA-seq data consisted of four batches, of which we only used one batch, the control digitonin-permeabilized cells, containing 11,868 cells. We kept cells in the TEA-seq data that had between 500 and 2,750 unique genes, at least 1000 unique peaks, and were not part of a cluster that expressed the mouse control protein. We kept DOGMA-seq data that had at least 500 peak counts, between 200 and 30,000 gene counts, 20 and 10,000 epitope counts, and fewer than 50 rat control epitopes.

In the TEA-seq integration experiment, we used Harmony-corrected PCA, LSI, and centered log ratio (CLR) transformed^4^,^80^ counts were used as the embeddings for the scRNA-seq, scATAC-seq, and epitope data, respectively. In the DOGMA-seq re-pairing experiment, we used scVI, PeakVI, and Harmony-corrected scaled CLR counts as input to scPairing. During pairing, we used similarity cutoff *ε* = 0.1.

The joint scRNA-seq and scATAC-seq dataset used to generate a new trimodal dataset was the NeurIPS 2021 benchmarking data. The epitope data came from the same NeurIPS 2021 benchmarking data,^38^ except from the CITE-seq data. We discarded the scRNA-seq data from the CITE-seq data to retain only the epitope sequencing. For low-dimensional representations, we used scVI and PeakVI for the scRNA-seq and scATAC-seq data, respectively. For the epitopes, we used Harmony-corrected scaled CLR transformed counts. During pairing, we used similarity cutoff *ε* = 0.1.

## Data availability

The 10x Multiome and CITE-seq BMMCs from the Multimodal SingleCell Data Integration Challenge at NeurIPS 2021 is available via GEO accession GSE194122.^38^ The SHARE-seq mouse skin dataset is available via GEO accession GSE140203.^2^ The joint scRNA-seq and scATAC-seq human cerebral cortex data is available via GEO accession GSE204682.^42^ The human PBMC CITE-seq data is available via GEO accession GSE164378 and the New York Genome Center.^14^

The pre-trained model checkpoints for CellPLM and scGPT were downloaded from the authors’ sources on Dropbox ^81^ and Google Drive,^82^ respectively.

The scRNA-seq raw counts data from the Wang *et al*. paired retina dataset is available via the UCSC Cell Browser^83^ dataset ID chang-retina-atlas and the scATAC-seq fragment files are available via GEO accession GSE196235.^44^ The snRNA-seq raw counts data and snATAC-seq fragment files from the Liang *et al*. unpaired retina dataset is available via the UCSC Cell Browser^83^ dataset ID retina-atac.^43^

The paired Ulezko Antonova *et al*. bridge dataset of human tonsil cells is available via GEO accession GSE247692.^49^ The paired and unpaired human tonsil cell atlas is available via Zenodo.^51^,^84^The paired human intestine dataset is available via HuBMAP ID HBM692.JRZB.356, while the data matrices and fragment files are available through Data Dryad via DOI 10.5061/dryad.0zpc8672f.^52^

The paired Hu *et al*. bridge dataset of ccRCC cells is available via Zenodo ^57^,^85^ and the paired 10x PBMCs bridge dataset is available from 10x Genomics.^58^ The unpaired ccRCC scATAC-seq and scRNA-seq data are available via GEO accession GSE207493.^56^

The TEA-seq dataset is available via GEO accession GSE158013.^25^ The DOGMA-seq dataset is available via GEO accession GSE156478.^24^

## Code availability

The scPairing model is available on GitHub (https://github.com/Ding-Group/scPairing). The scripts and computational notebooks used to produce and analyze the results are deposited in Zenodo (https://doi.org/10.5281/zenodo.13910219).

## Acknowledgements

We thank Minuk Ma for his comments on the manuscript.

This work was supported by a Discovery grant from the Natural Sciences and Engineering Research Council (NSERC) of Canada, and a department startup fund from the University of British Columbia (to J.D.). J.D. is a Canada Research Chair and is supported by the Canadian Institutes of Health Research through the Canada Research Chair Program. The computational resource is partially supported by the Canada Foundation for Innovation & John. R. Evans Leader Fund (to J.D.). This research was supported in part through the computational resources and services provided by Advanced Research Computing at the University of British Columbia. J.N. is supported by a Master’s scholarship from NSERC and a graduate scholarship from the province of British Columbia.

## Author’s contributions

J.N. and J.D. conceived the study. J.N. designed and wrote the code for scPairing. J.N. and C.V.R. ran computational experiments and interpreted the results. J.D. supervised the project. J.N., C.V.R., and J.D. approved the final manuscript.

## Supplementary Figures

**Figure S1:**
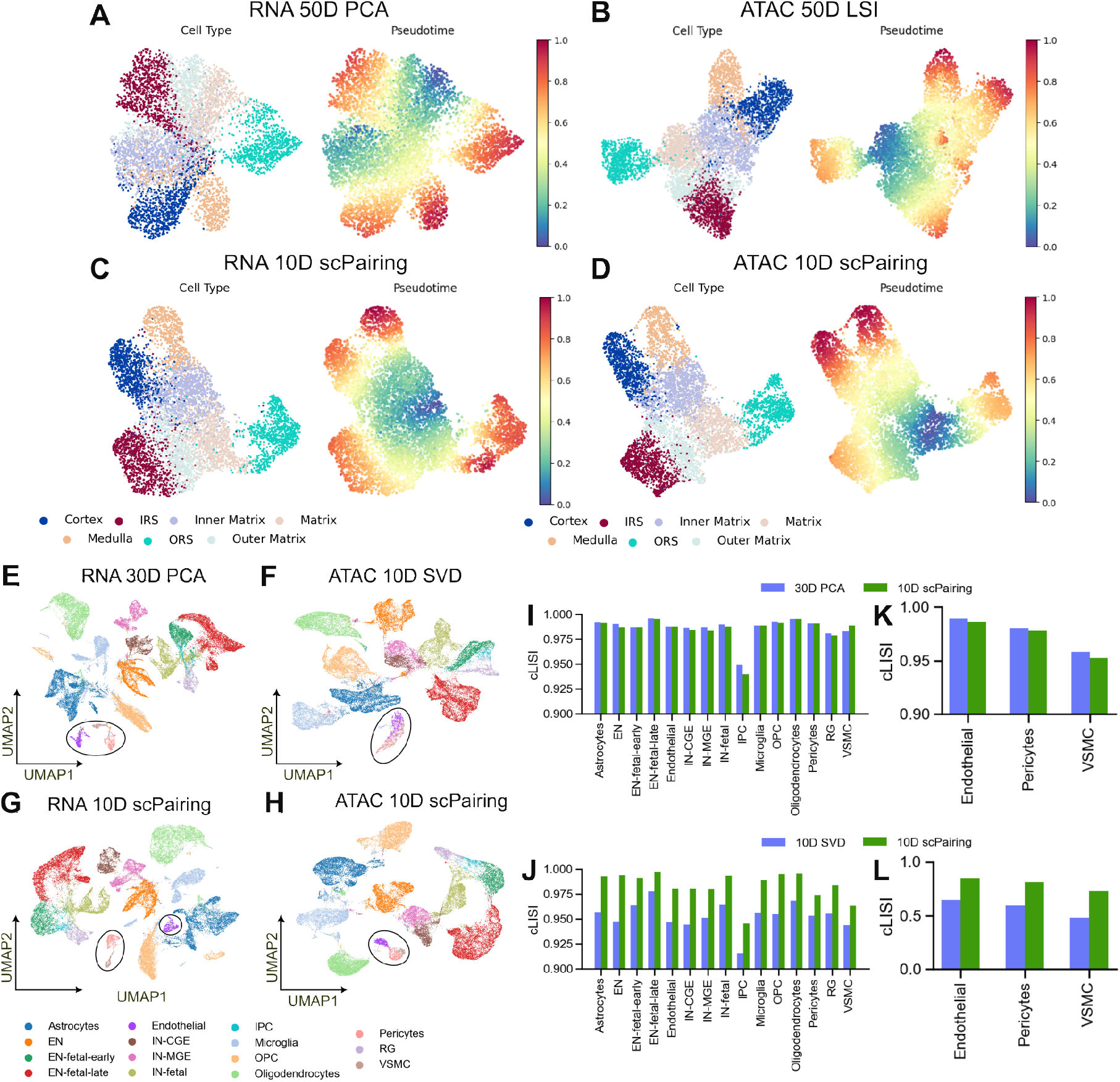
Alignment of modalities transfers modality-specific information onto both modalities, related to Fig. 2. (A–D) UMAP visualization of cell type and pseudotime of mouse skin data using scRNA-seq PCA embeddings (A), scATAC-seq LSI embeddings (B), scPairing’s transformation of the PCA embeddings (C), and scPairing’s transformation of the LSI embeddings (D). (E–L) Alignment of human cerebral cortex multiomics data. (E and F) UMAP visualization of cell types following the data processing procedure from Zhu *et al*., ^42^ with the 30-dimension PCA embeddings (E) and the 10-dimension SVD embeddings (F). (G and H) UMAP visualization of cell types after applying scPairing to the PCA and SVD embeddings from (E) and (F). The three blood vessel subtypes are highlighted in the black circle. (I and J) Comparison of cell type LISI (cLISI) between the PCA embeddings and scPairing embeddings in the scRNA-seq modality (I), and between the SVD embeddings and scPairing embeddings in the scATAC-seq modality (J). (K and L) Comparison of cLISI computed only on the three blood vessel cell subtypes. (K) compares the cLISI between the PCA embeddings and scPairing embeddings in the scRNA-seq modality. (L) compares the cLISI between the SVD embeddings and scPairing embeddings in the scATAC-seq modality.

**Figure S2:**
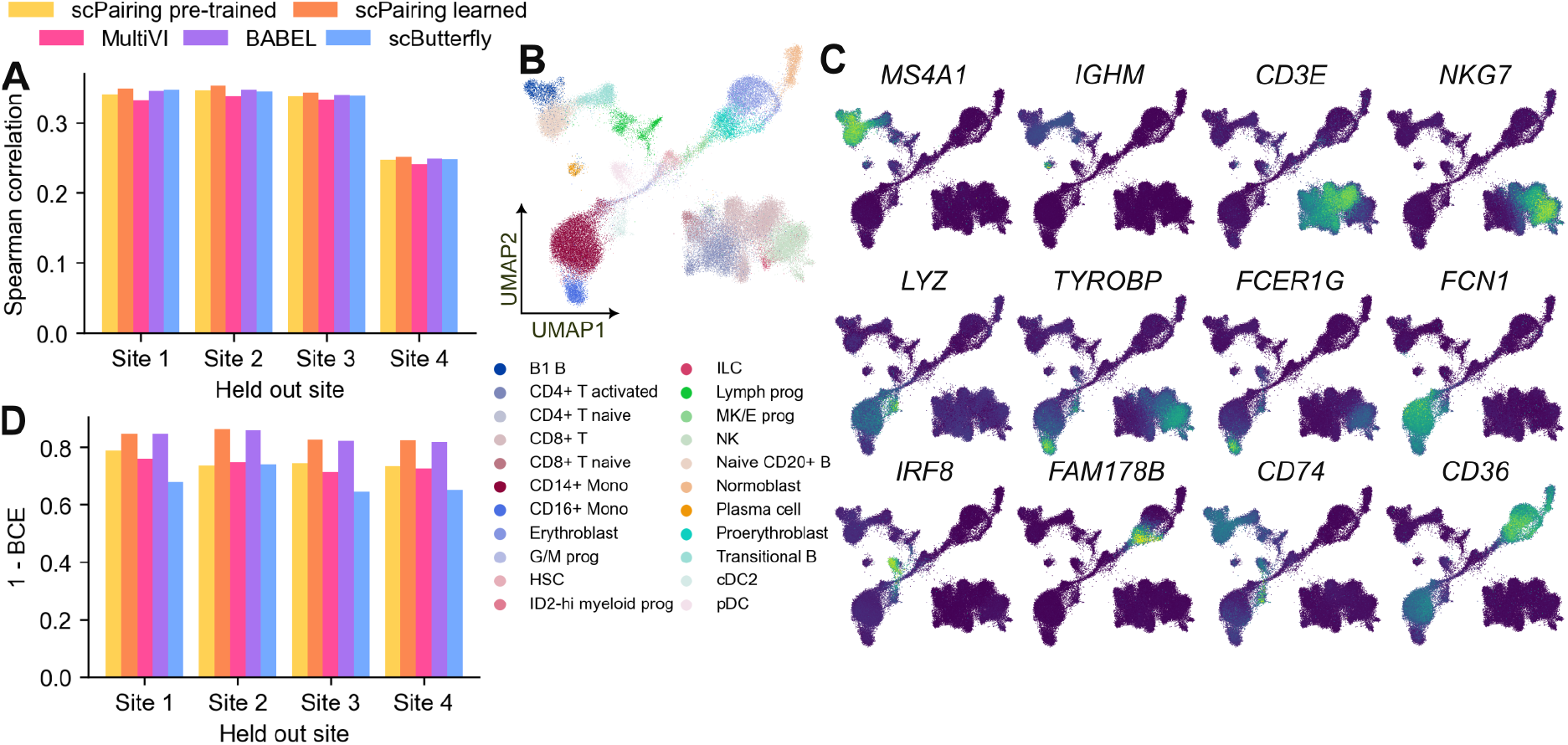
Benchmarking cross-modal imputation using pre-trained and learned decoders, related to Fig. 2. (A) Comparison of cross-site scATAC-seq to scRNA-seq imputation with Spearman correlations between imputed and true gene counts. (B) UMAP visualization of the scRNA-seq BMMC benchmarking dataset using scVI embeddings. (C) Visualization of scPairing marker gene expression imputations. We applied scVI and PeakVI pre-trained decoders to impute the counts. (D) Comparison of cross-site scRNA-seq to scATAC-seq imputation with binary cross-entropy between the imputed and true binarized peaks.

**Figure S3:**
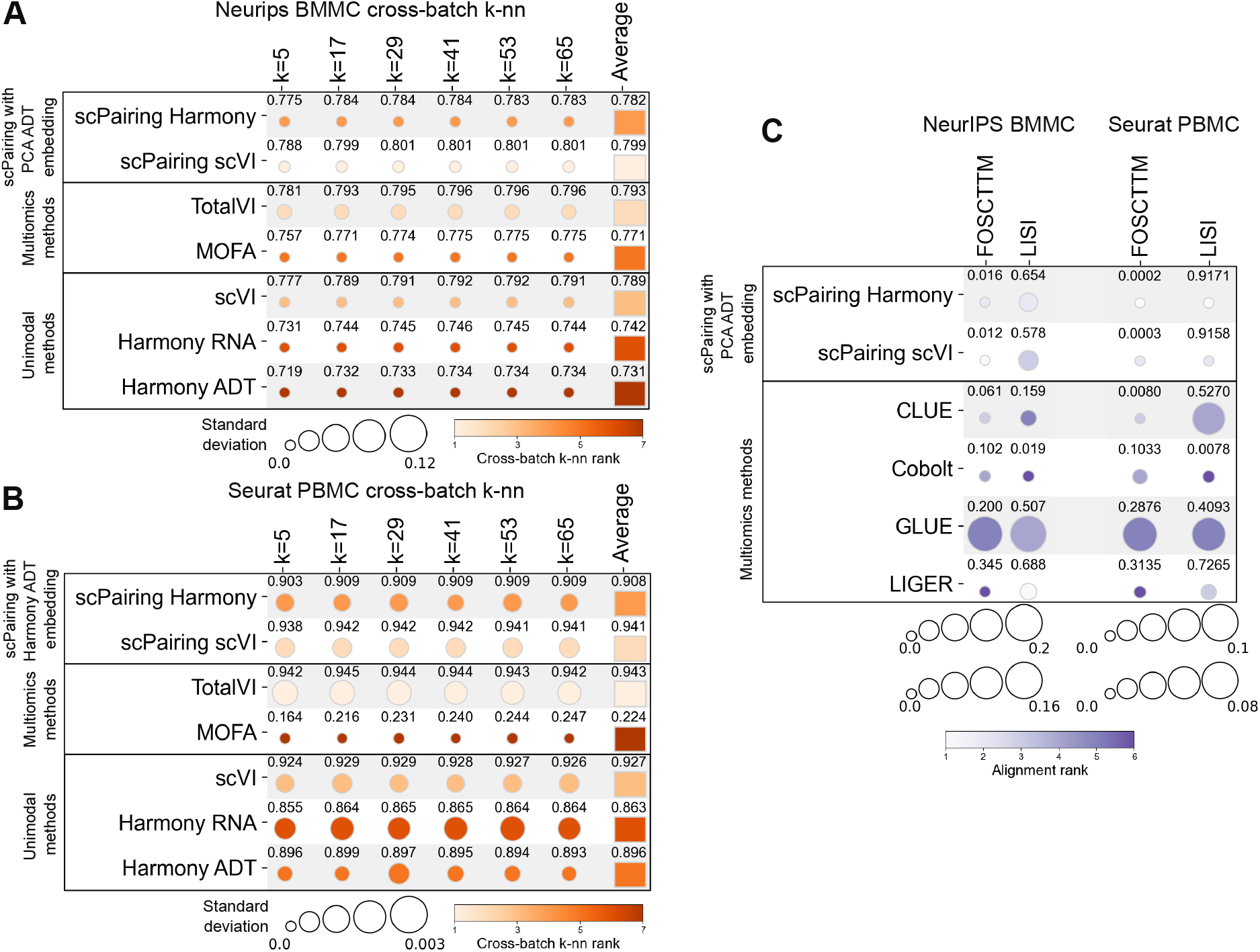
Benchmarking integration of CITE-seq data. (A and B) Comparison of biological structure preservation in the BMMC data (A) and the PBMC data (B) quantified by *k*-nearest neighbor cell type classification accuracies, where we predicted cell types for one batch given the cells from the remaining batches (cross-batch *k*-nn) or predicted cell types for held-out cells using 10-fold cross-validation (10-fold *k*-nn). (C) Comparison of modality alignment performance with FOSCTTM and LISI computed across all cells in the BMMC data (left two columns) and the PBMC data (right two columns). scPairing Harmony and scPairing scVI refers to scPairing with Harmony-corrected PCA and scVI used as scRNA-seq embedding, respectively. All experiments were repeated for five trials. The mean of each metric is labeled and the standard deviation is given by the circle size.

**Figure S4:**
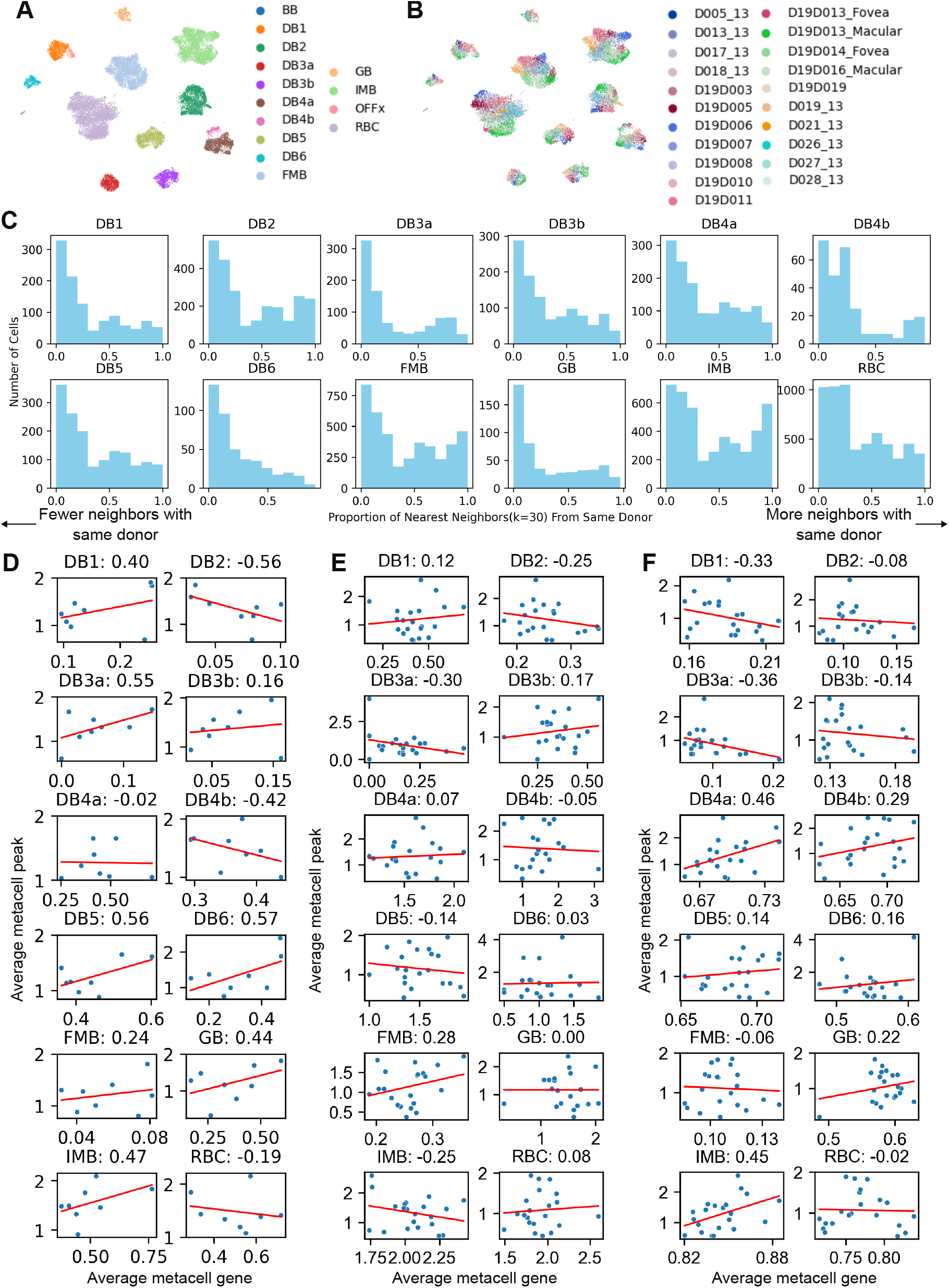
Donor-aggregated metacells in retina data, related to Fig. 3 and Methods. (A and B) UMAP visualization of bipolar retina cells from Wang *et al*. ^44^ *using LSI dimensionality reduction without batch correction, colored by cell subtype (A) and donor (B). (C) Proportion of nearest neighbors (k* = 30) having the same donor for each cell subtype. (D) Correlations computed on the Wang *et al*. paired multiomics dataset using donor-aggregated metacells. The data contained four donors with two retina each, for eight batches in total. (E) Correlations computed on the artificial pairings using donor-aggregated metacells. The data contained 21 donors. (F) Correlations computed on the gene activities and CCA imputations using donor-aggregated metacells. This is the same data as in (E), with 21 donors.

**Figure S5:**
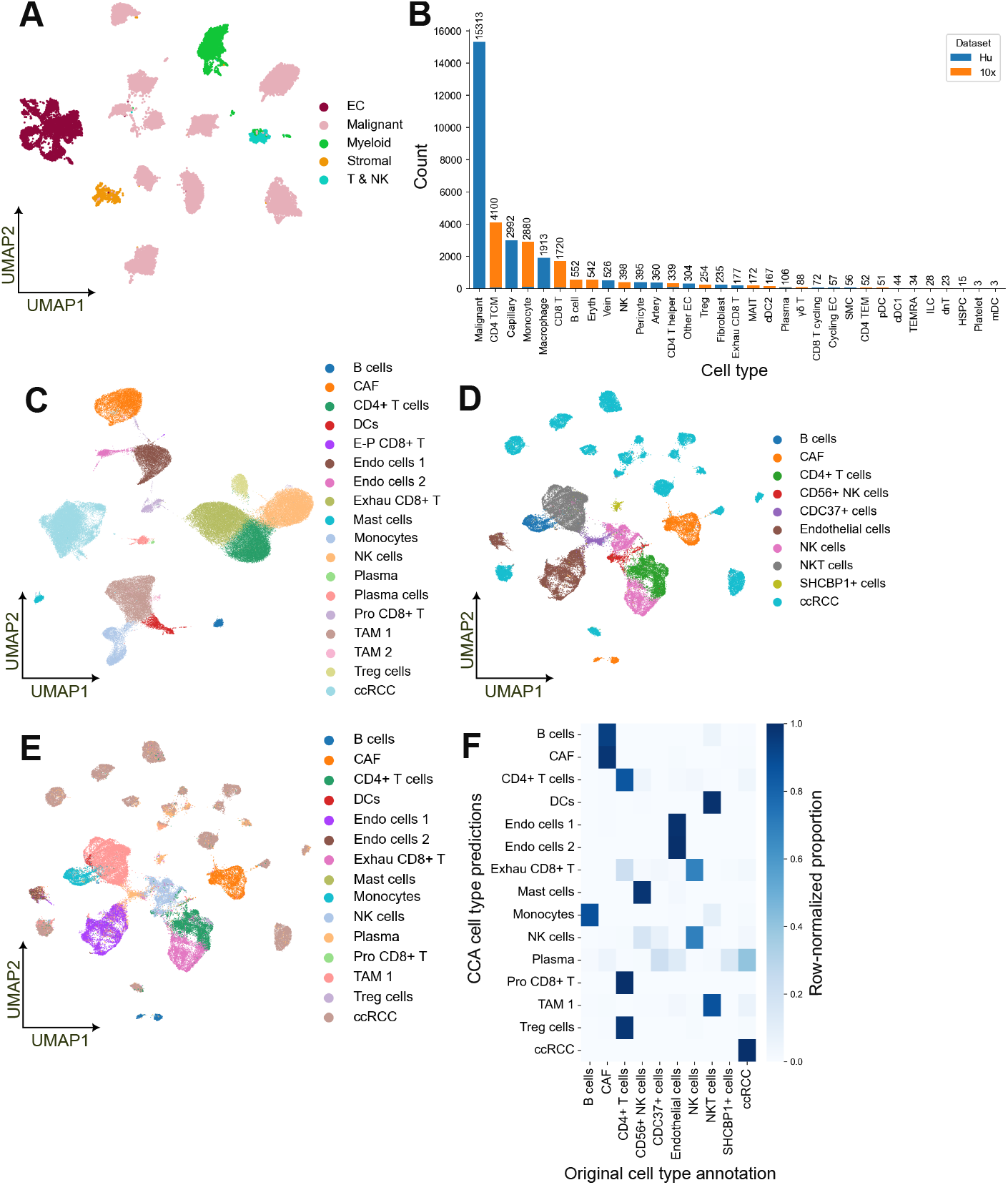
Cell type annotations of ccRCC bridge and single-modality data, related to Fig. 6. (A) UMAP visualization of cell types in the Hu *et al*. data. ^57^ (B) Cell type counts in the bridge data after adding PBMCs from 10x Multiome. ^58^ (C) Original cell type annotations of ccRCC scRNA-seq data from Yu *et al*. ^56^, *whose annotations were transferred onto the scATAC-seq data in (D and E). (D) Original cell type annotations of the ccRCC scATAC-seq data from Yu et al*., which were annotated on the basis of gene scores, differentially expressed peaks, and transcription factor analysis. ^56^ (E) Cell type annotations of the ccRCC scATAC-seq data in (D) following label transfer using CCA. (F) Comparison of CCA cell type annotations of the scATAC-seq data in (C) against the original annotations in (D). The heatmap is normalized per row.

**Figure S6:**
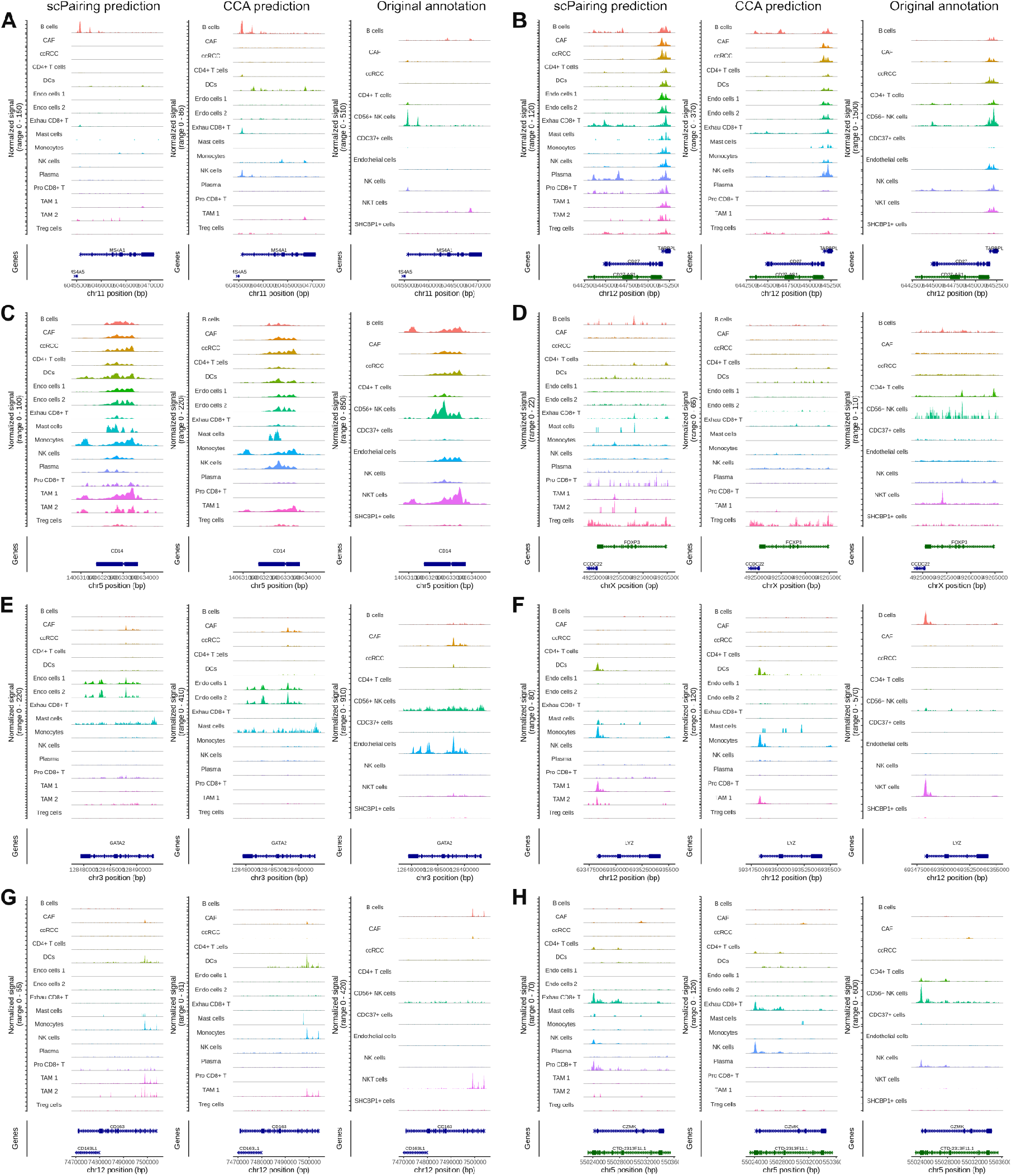
Accessibility profiles of ccRCC scATAC-seq profiles at marker genes, related to Fig. 6. For each gene, the accessibilities are grouped by cell type annotations from scPairing (left column), CCA (middle column), and the original annotations (right column). (A–H) Accessibility plots for *MS4A1* (A, a B cell marker), *CD27* (B,), *CD14* (C, expressed in monocytes), *FOXP3* (D, a regulatory T cell marker), *GATA2* (E, mast cells), *LYZ* (F, monocytes/macrophages/dendritic cells), *CD163* (G, monocytes/macrophages/dendritic cells), and *GZMK* (H, exhausted CD8^+^ T cells). Accessibility for E-P CD8+ T cells are omitted due to low cell number. 8

**Figure S7:**
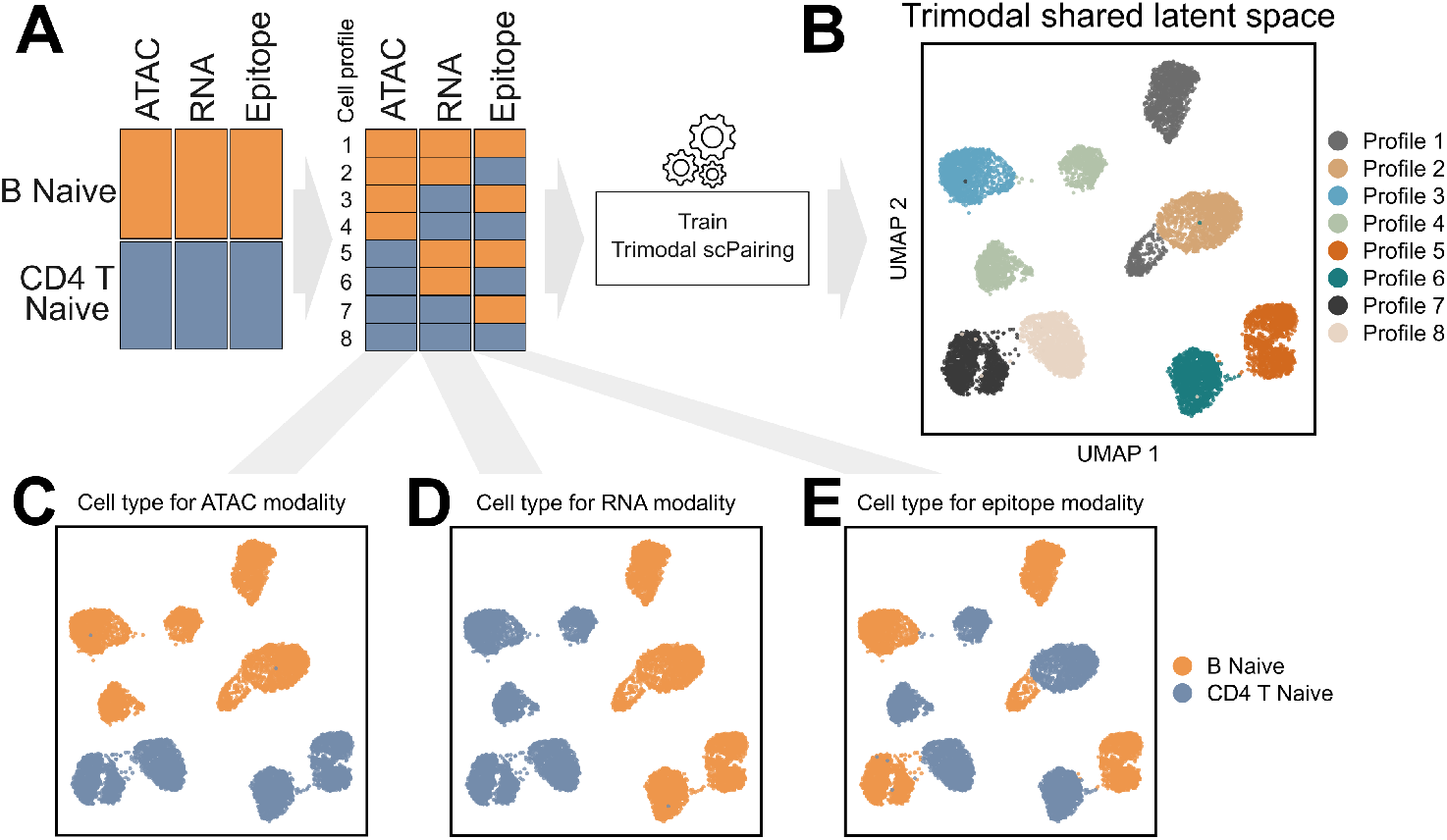
scPairing separates cells with shuffled modalities. (A) Schematic of the modality shuffling, with eight profiles generated from all combinations of scATAC-seq, scRNA-seq, and epitope modalities of B naive and CD4 T naive cells. (B) UMAP visualization of scPairing embeddings of the eight shuffled profiles. (C–E) UMAP visualization of the individual modalities.

## Supplementary notes

### S1 Trimodal pairing algorithm

#### Algorithm 1

Trimodal pairing

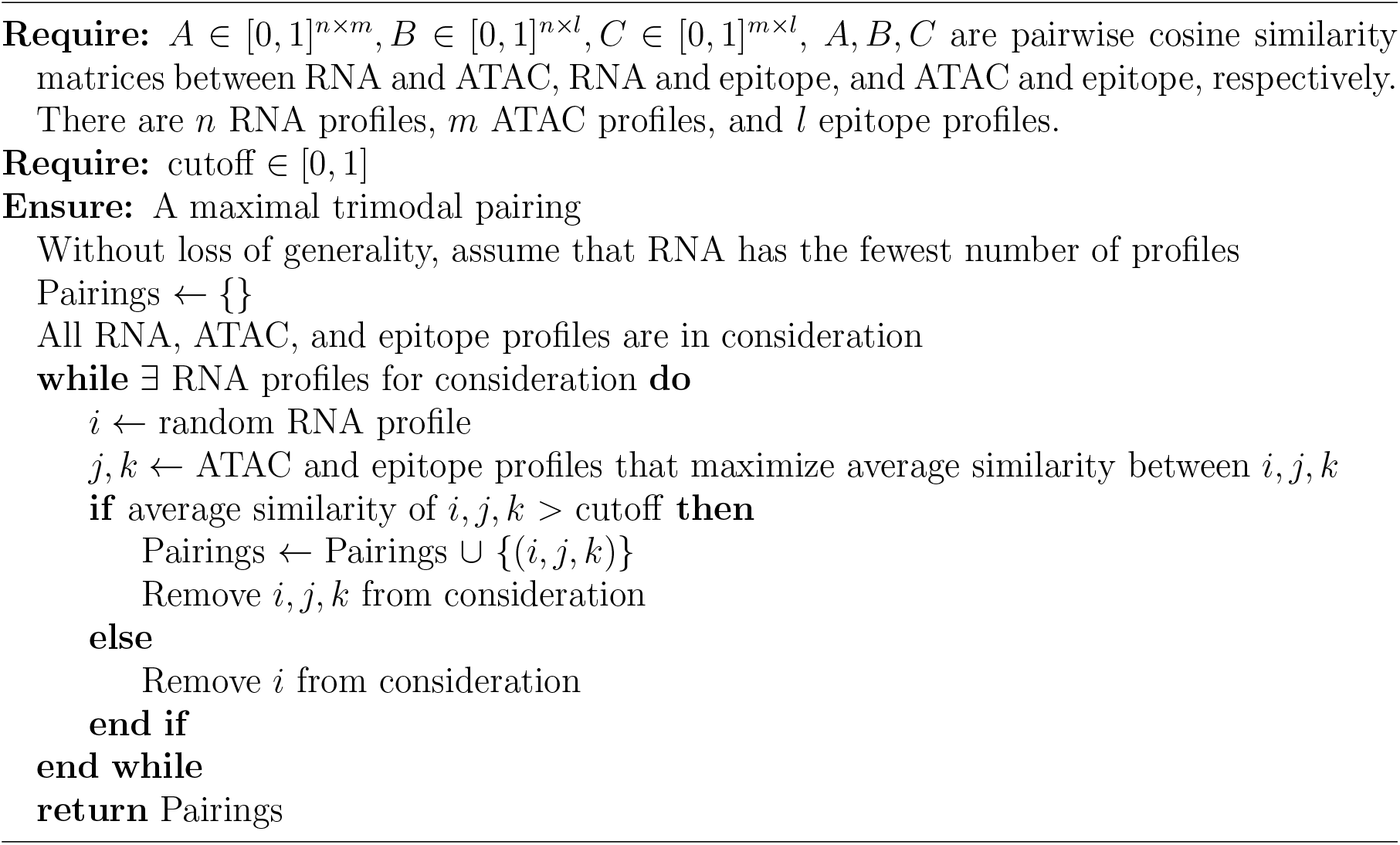

## Supplementary tables

**Table S1:**
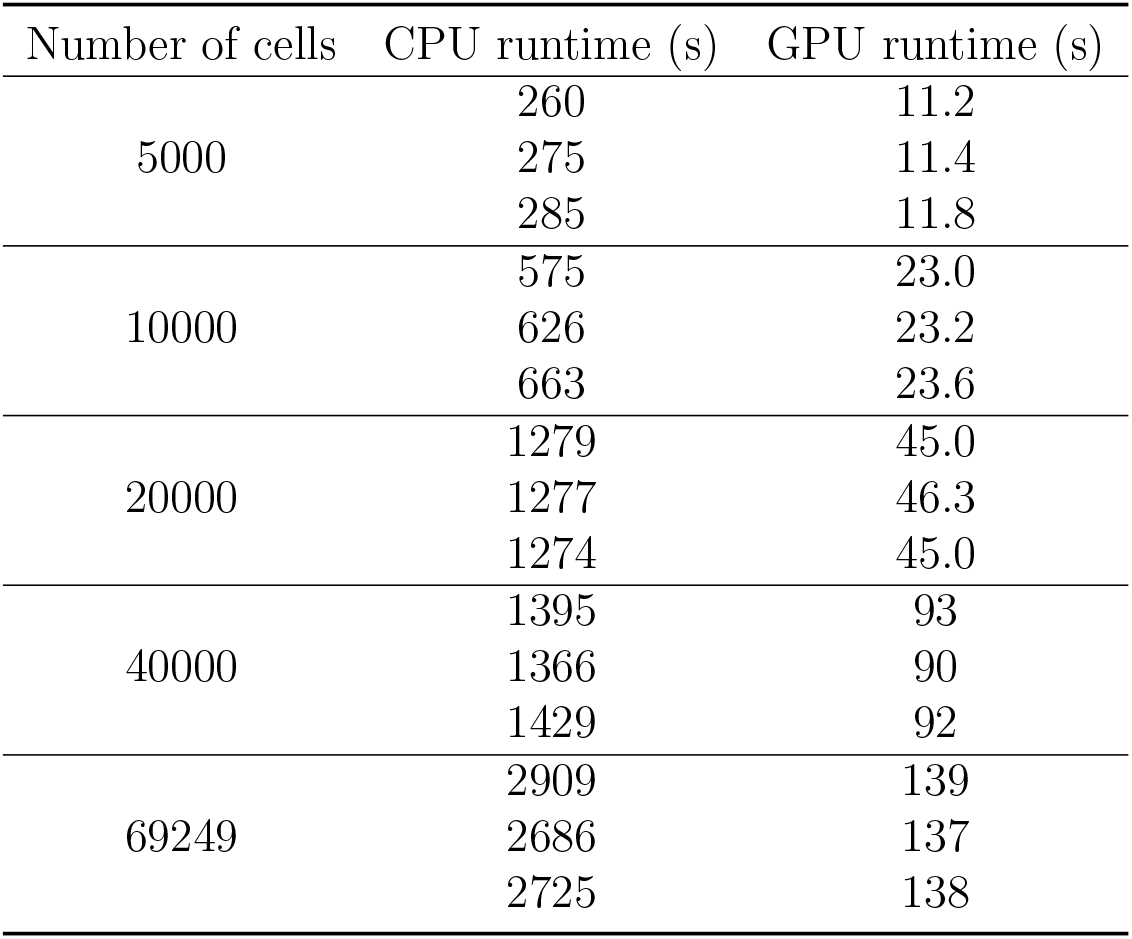
Runtime of scPairing. Runtime using a single Intel Xeon Gold 6130 CPU or a single NVIDIA Tesla V100 GPU in seconds across five dataset sizes. Each dataset size was run for three trials. The result of each trial is reported.

| Number of cells | CPU runtime (s) | GPU runtime (s) |
| --- | --- | --- |
| 5000 | 260 | 11.2 |
|  | 275 | 11.4 |
|  | 285 | 11.8 |
| 10000 | 575 | 23.0 |
|  | 626 | 23.2 |
|  | 663 | 23.6 |
| 20000 | 1279 | 45.0 |
|  | 1277 | 46.3 |
|  | 1274 | 45.0 |
| 40000 | 1395 | 93 |
|  | 1366 | 90 |
|  | 1429 | 92 |
| 69249 | 2909 | 139 |
|  | 2686 | 137 |
|  | 2725 | 138 |

**Table S2:** Average FOSCTTM and proportion of cells paired on eight variations of scPairing losses, related to Fig. 4. Each of the eight combinations of modality discriminative (MD) loss, batch discriminative (BD) loss, and cosine similarity (CS) loss were tested. The FOSCTTMs and proportion of cells paired were calculated from the test data after applying each scPairing variant. The proportion of cells paired is the number of pairings produced by the linear sum assignment algorithm divided by the total number of cells in the test BMMC data, or the number of pairings divided by the total number of snATAC-seq cells in the retina data. Each scPairing variant was evaluated across five trials, with the mean and standard error reported.

| MD loss | BD loss | CS loss | Test BMMC data |  | Retina data |
| --- | --- | --- | --- | --- | --- |
|  |  |  | FOSCTTM | Proportion of cells paired | Proportion of cells paired |
| ✓ | ✓ | ✓ | 0.010442<br>± 0.000151 | 0.991370<br>± 0.000695 | 0.845802<br>± 0.015269 |
| ✓ | ✓ | ✗ | 0.010712<br>± 0.000213 | 0.943339<br>± 0.004580 | 0.404926<br>± 0.128817 |
| ✓ | ✗ | ✓ | 0.010481<br>± 0.000112 | 0.992948<br>± 0.000384 | 0.762668<br>± 0.004222 |
| ✓ | ✗ | ✗ | 0.010654<br>± 0.000120 | 0.946065<br>± 0.002797 | 0.272155<br>± 0.076137 |
| ✗ | ✓ | ✓ | 0.010582<br>± 0.000322 | 0.990906<br>± 0.000549 | 0.808498<br>± 0.015745 |
| ✗ | ✓ | ✗ | 0.010769<br>± 0.000327 | 0.935046<br>± 0.003350 | 0.299533<br>± 0.098023 |
| ✗ | ✗ | ✓ | 0.010537<br>± 0.000187 | 0.993273<br>± 0.000207 | 0.769556<br>± 0.004856 |
| ✗ | ✗ | ✗ | 0.010660<br>± 0.000131 | 0.945184<br>± 0.001766 | 0.348049<br>± 0.075139 |

**Table S3:**
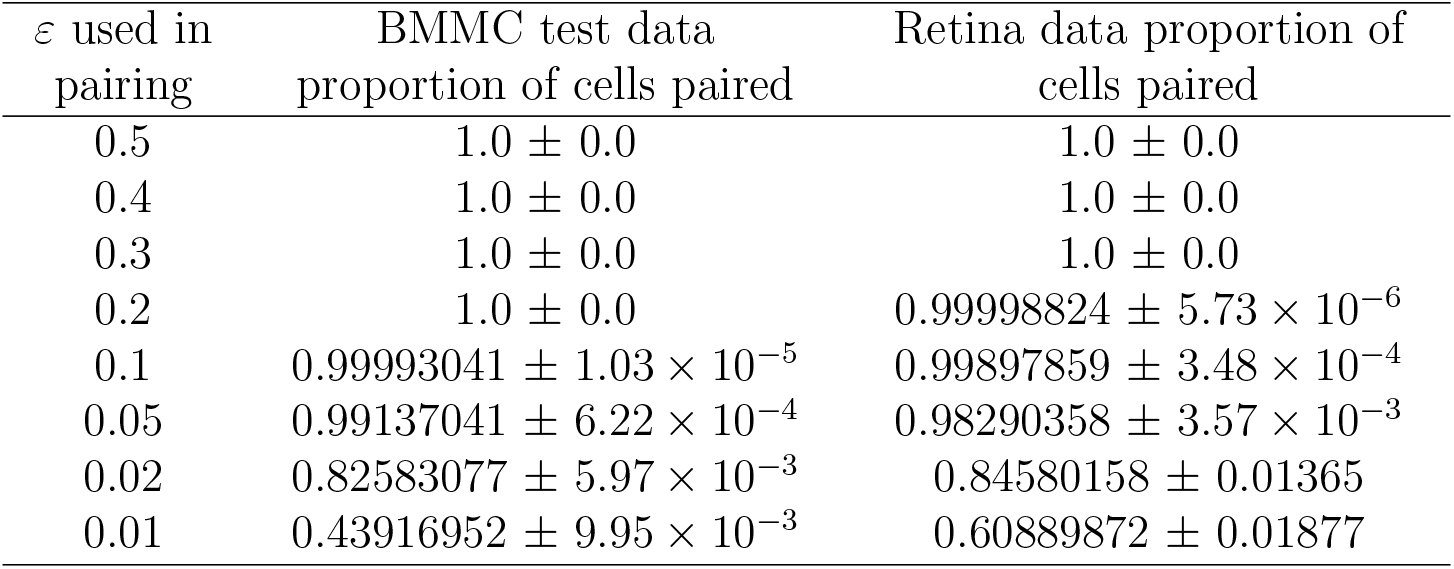
Average proportion of cells paired when varying *ε*, related to Fig. 4. The proportion of cells paired is the number of pairings produced by the linear sum assignment algorithm divided by the total number of cells in the case of the BMMC test data, or the total number of scATAC-seq cells in the retina data. Each scPairing variant was evaluated across five trials, with the mean and standard error reported.

**Table S4:**
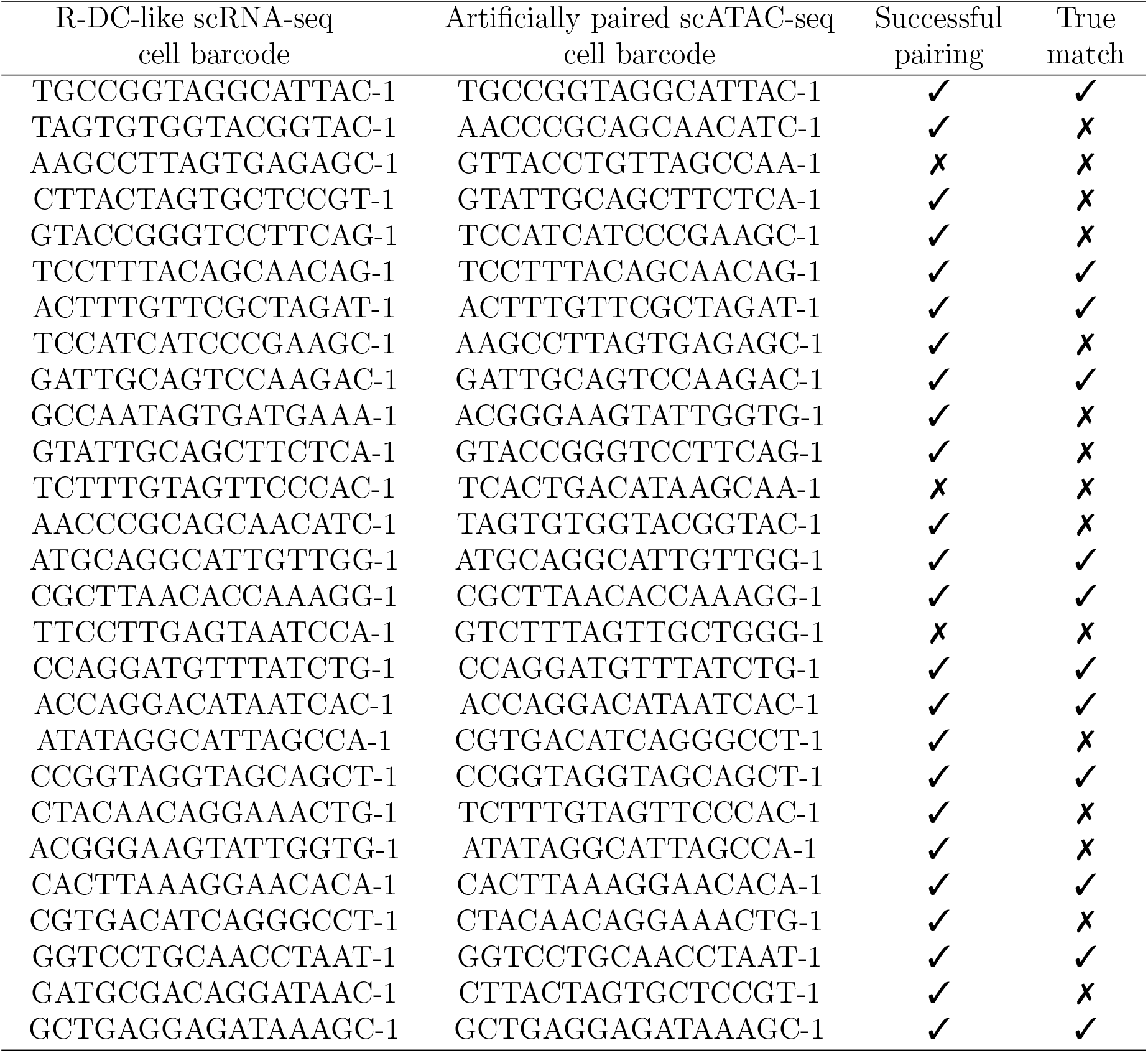
Re-pairings of human tonsil R-DC-like cells, related to Fig. 5. The cells in the first column were identified as likely R-DC-like cells from their gene expression profiles. The second column indicates the scATAC-seq profile that was paired with each R-DC-like gene expression profile. The third column indicates whether the scRNA-seq and scATAC-seq profiles both belonged to R-DC-like cells. The fourth column indicates whether the scRNA-seq and scATAC-seq profiles came from the same cell, meaning scPairing recapitulated the true pairing of the two modalities. The cell barcodes listed have had their prefixes stripped off, but no duplicate barcodes are present in the R-DC-like cells.

**Table S5:**
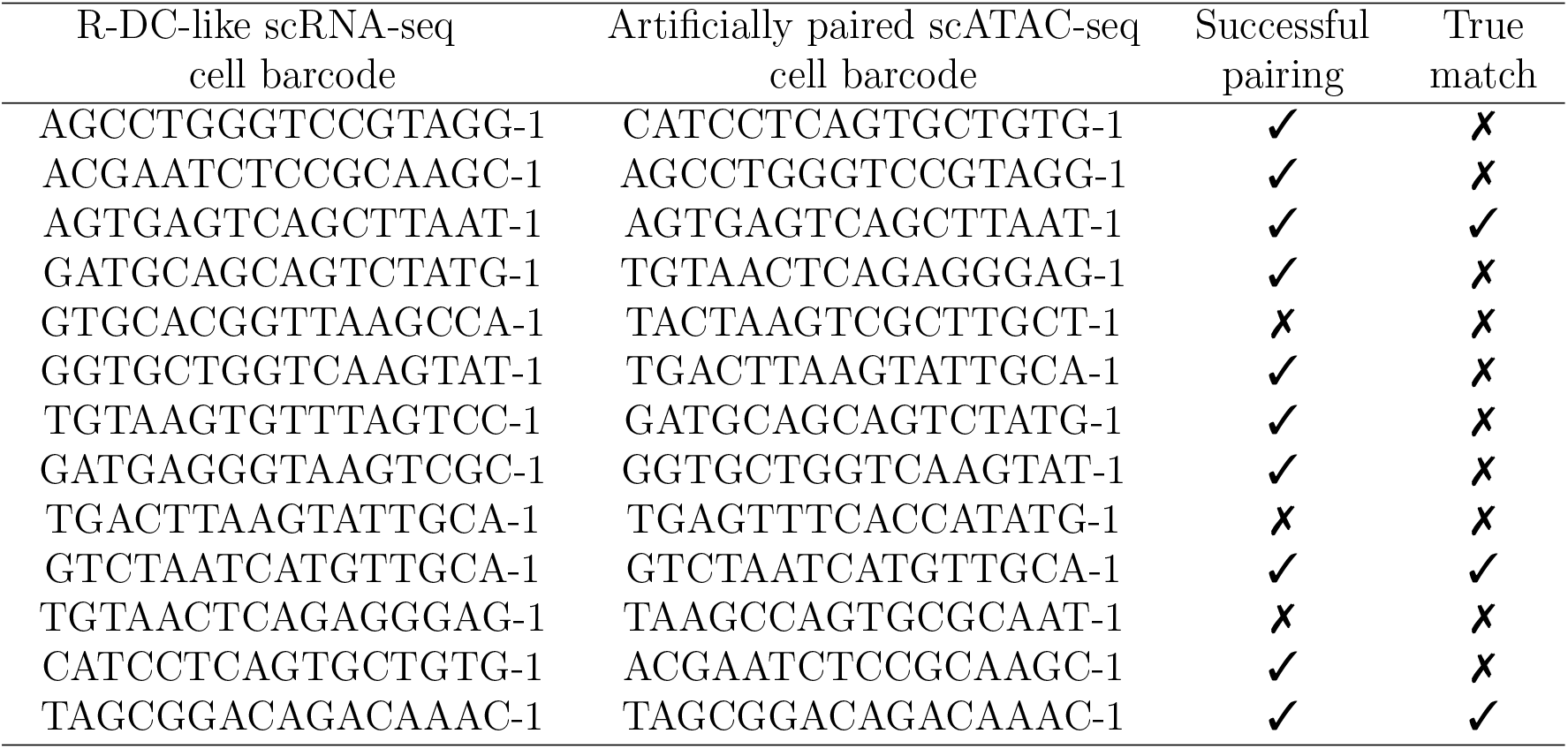
Re-pairings of intestine R-DC-like cells, related to Fig. 5. The cells in the first column were identified as likely R-DC-like cells from their gene expression profile. The second column indicates the scATAC-seq profile that was paired with each R-DC-like gene expression profile. The third column indicates whether the scRNA-seq and scATAC-seq profiles both belonged to R-DC-like cells. The fourth column indicates whether the scRNA-seq and scATAC-seq profiles came from the same cell, meaning scPairing recapitulated the true pairing of the two modalities. The cell barcodes listed have had their prefixes stripped off, but no duplicate barcodes are present in the R-DC-like cells.

**Table S6:** Mean and median pairwise modality cosine similarities, and FOSCTTM between the true matching triplets from DOGMA-seq, related to Fig. 7. The cosine similarities and FOSCTTM are calculated using the embeddings produced by applying scPairing with TEA-seq as the bridge data. The similarity and FOSCTTM is computed between each pair of modalities.

|  | RNA-ATAC | RNA-epitope | ATAC-epitope |
| --- | --- | --- | --- |
| Mean Cosine Similarity | 0.9579 | 0.9060 | 0.9253 |
| Median Cosine Similarity | 0.9640 | 0.9183 | 0.9366 |
| FOSCTTM | 0.05378 | 0.1620 | 0.1294 |

